# Exploring the bounded rationality in human decision anomalies through an assemblable computational framework

**DOI:** 10.1101/2023.10.16.562648

**Authors:** Yi-Long Lu, Yang-Fan Lu, Xiangjuan Ren, Hang Zhang

## Abstract

Some seemingly irrational decision behaviors (anomalies), once seen as flaws in human cognition, have recently received explanations from a rational perspective. The basic idea is that the brain has limited cognitive resources to process the quantities (e.g., value, probability, time, etc.) required for decision making, with specific biases arising as byproducts of the resource allocation that is optimized for the environment. While appealing for providing normative accounts, the existing resource-rational models have limitations such as inconsistent assumptions across models, a focus on single environmental factors, and limited coverage of decision anomalies. One challenging anomaly is the peanuts effect, a pervasive phenomenon in decision-making under risk that implies an interdependence between the processing of value and probability. To extend the resource rationality approach to explain the peanuts effect, here we develop a computational framework—the Assemblable Resource-Rational Modules (ARRM)—that integrates ideas from different lines of boundedly-rational decision models as freely assembled modules. The framework can accommodate the joint functioning of multiple environmental factors, and allow new models to be built and tested along with the existing ones, potentially opening a wider range of decision phenomena to bounded rationality modeling. For one new and three published datasets that cover two different task paradigms and both the gain and loss domains, our boundedly-rational models reproduce two characteristic features of the peanuts effect and outperform previous models in fitting human decision behaviors.

## Introduction

Many decision problems in the real world can be framed as choice among gambles whose outcomes are probabilistic. What principles do human decisions follow? An early influential theory was the Expected Utility Theory (EUT; von Neumann & Morgenstern, 1944), which assumes that humans maximize expected utility, as any rational agents would do. However, following the pioneering experimental work of Allais (1953), significant violations of EUT were identified in human decisions. These anomalies motivated Kahneman and Tversky’s Cumulative Prospect Theory (CPT; Kahneman & Tversky, 1979; Tversky & Kahneman, 1992), which retains EUT’s assumption of expected utility maximization but introduces a distortion of probability in the computation of expected utility, typically an overestimation of small probability and underestimation of large probability (i.e., an inverted-*S*-shaped distortion curve). With probability distortion modeled by a one- or two-parameter function (Gonzalez & Wu, 1999; Kahneman & Tversky, 1979; Prelec, 1998), CPT provides a good fit to a variety of human decision behaviors, including those that challenge EUT, and thus becomes one of most widely used decision models in economics, psychology, and neuroscience (Glimcher & Fehr, 2014).

From a cognitive view, CPT is unsatisfying in its lack of mechanistic explanation for the observed value or probability distortions. Even as a descriptive model for human behaviors, it has some critical limitations. For example, it assumes a separability of value and probability, that is, neither does the utility function depend on probability, nor does the probability distortion function depend on value. This separability assumption turns out to be oversimplified, failing to capture the rich interaction of value and probability in human decisions (Peterson et al., 2021). A well-documented phenomenon is known as the “peanuts effect”: People tend to take more risk when the value at stake is small (Estle et al., 2006; Fehr-Duda et al., 2010; Green et al., 1999; Jones & Oaksford, 2011; Kachelmeier & Shehata, 1992; Prelec & Loewenstein, 1991; Weber & Chapman, 2005).

These highly patterned violations of EUT and even CPT lead some researchers to consider an entirely different possibility: people may not compute or maximize expected utility at all, but instead rely on simple heuristic algorithms to make decisions (Gigerenzer et al., 2011), often discarding part of the available information. For example, the maximax heuristic compares the best outcome of each gamble and chooses the one whose best outcome is the best (Coombs et al., 1970); the similarity heuristic ignores the dimension that is similar for different gambles and focuses on the other dimension (Rubinstein, 1988). Numerous different heuristic decision models have been proposed (Birnbaum & LaCroix, 2008; Brandstätter et al., 2006; Leland, 1998). However, one problem with heuristic models is a lack of consensus on which heuristic to use in which situation (Callaway et al., 2022). An even more serious problem is that these heuristics may not produce the same types of decision biases as humans do (Birnbaum, 2008). Indeed, no known heuristic models have outperformed CPT in capturing the comprehensive patterns of human decision behaviors (Erev et al., 2010).

A third modeling approach for human decisions, alternative to CPT and heuristics, is bounded rationality (Anderson, 1990; Simon, 1955), where humans are modeled as agents who have limited computational power, but who use it efficiently to achieve adaptive behaviors, with decision biases arising as byproducts. An early model along this line is Decision by Sampling (Stewart et al., 2006), which explains the inverted-*S*-shaped probability distortion as the consequence of an adaptive representation of probability according to the event probabilities in the environment. More recently, such bounded rationality has been formulated within the framework of information theory (C. J. Bates et al., 2019; C. A. Sims, 2003; C. R. Sims, 2016), providing normative explanations for the subjective distortions of value (C. J. Bates et al., 2019; Gershman & Bhui, 2020; C. R. Sims, 2016), probability (Bhui & Gershman, 2018; Zhang et al., 2020), numerosity (Heng et al., 2020), and temporal delay (Gershman & Bhui, 2020). What lies at the heart of these models is the assumption of limited cognitive resources and a rational use of these resources. This approach, as a branch of bounded rationality, is thus called resource rationality (Bhui et al., 2021; Lewis et al., 2014; Lieder & Griffiths, 2020).

The resource rationality approach is appealing, because it can provide normative accounts for how human decisions may be determined by both the environmental and cognitive constraints. Meanwhile, it is faced with some challenges. First, most of its applications focus on rational resource allocation controlled by one single dimension of the environment, which could be an oversimplification of human decision problems. A full treatment needs to deal with the joint influence of multiple environmental factors. Second, it is common for different alternative assumptions to be used in the resource-rational modeling of different phenomena. Because most studies do not compare across alternative resource-rational models, it is often unknown whether a specific assumption is necessary for a specific phenomenon and which assumption best describes the optimalization problem posed on and solved by human cognition. Third, only a small portion of known decision anomalies have been addressed by the approach (Frydman & Jin, 2021; Gershman & Bhui, 2020; Summerfield & Parpart, 2022). It can hardly explain a wider range of decision phenomena unless its model space is significantly expanded.

The peanuts effect, briefly mentioned earlier, is a pervasive anomaly in decision-making under risk that has long challenged traditional explanations. As we will demonstrate later, this phenomenon implies an interdependence between the processing of value and probability, necessitating more powerful resource-rational models. Building on previous resource-rational work that models cognitive processing as information transmission through a constrained communication channel (Bhui et al., 2021; Dayan & Abbott, 2001; C. R. Sims, 2018; Wei & Stocker, 2015), we developed the Assemblable Resource-Rational Modules (ARRM) framework, which specifies each module of resource-rational models (e.g., cognitive resources, prior in memory, encoding and decoding schemes) and provides a taxonomy to classify their variants. This framework enables us to formulate the joint effects of multiple environmental factors on resource allocation and to assemble assumptions from different previous models into new resource-rational models. To test which assumptions best explain the peanuts effect, we applied ARRM to decision under risk to construct 10 new models and compared their goodness-of-fit to human data in one new and three published datasets spanning two task paradigms and both gain and loss domains. The resource-rational model that best explained the peanuts effect was the bounded log-odds model (Zhang et al., 2020) enhanced with the assumptions of rational inattention (Gershman & Bhui, 2020; C. A. Sims, 2003) and structural prior (Pleskac & Hertwig, 2014), which provides a complete description of human risky decision behaviors including two characteristic features of the peanuts effect. The model also outperformed a representative set of traditional decision models. We close with a discussion of ARRM’s other potential applications and its relationship to heuristic-based or attention-dependent decision models.

## The peanuts effect in decision under risk

Before diving into the resource-rational modeling of the peanuts effect, we briefly review the history of the anomaly and why it poses a challenge to almost all previous decision models. The peanuts effect was first presented by Markowitz (1952) as an (informal) observation that people may reverse their preferences between a sure reward and a gamble when the rewards of the two are proportionally enlarged. The set of choices constructed by Markowitz have the common format “X with certainty or one chance in ten of getting Y?”^1^, where Y=10X and X can be 10 cents, $1, $10, $100, $1,000, or $1,000,000. In all these choice pairs, the sure reward and the gamble have the same expected gain. The puzzling phenomenon is that most people seem to prefer the gamble for small X and Y but the sure reward for large X and Y. It receives the name “peanuts effect” in Prelec and Loewenstein (1991), defined as an elevated risk aversion for large gambles. The phenomenon has been repeatedly replicated in later studies on decision under risk (Binswanger, 1981; Estle et al., 2006; Fehr-Duda et al., 2010; Kachelmeier & Shehata, 1992; Weber & Chapman, 2005), for rating as well as choice tasks (Weber & Chapman, 2005), for real as well as virtual monetary rewards (Binswanger, 1981; Fehr-Duda et al., 2010; Kachelmeier & Shehata, 1992). As we will soon see, this phenomenon also widely exists in previous datasets that were not intended to test it, often unnoticed in the original publications.

### The theoretical difficulty in explaining the peanuts effect

Although the peanuts effect has been documented for decades, it has no satisfactory explanation yet. Below we briefly describe the theoretical explanations that have been proposed for the peanuts effect as well as decision theories that seem to have potential to explain it.

#### Convex-concave utility functions

In his original paper, Markowitz (1952) attributes the peanuts effect to utility functions that are convex in small values but concave in large values. According to such utility functions, the higher value of a small gamble is much more appealing than its paired sure reward, while for large gambles the appeal is weaker. However, there is little empirical evidence for such utility functions; what are widely accepted (Tversky & Kahneman, 1992) and agree with the non-parametric measurement (Gonzalez & Wu, 1999) of human decisions are power utility functions. Fehr-Duda et al. (2010) examined human choices for gambles with a range of different probabilities and argued that no utility functions may fully explain the peanuts effect.

The peanuts effect poses an anomaly not only for the expected utility theory, but also for CPT (Tversky & Kahneman, 1992) and many other theories (Prelec & Loewenstein, 1991; Wakker, 2010), because it challenges the *separability assumption* that value and probability are independently processed before their integration, that is, the utility function *u*(*x*) is determined only by value, and the probability weighting function *w*(*p*) only by probability. If such separability held, scaling up the values proportionally for the sure reward and gambles should not have influenced human preference between the two, at least under the widely accepted power utility function. In our Study 1, we will present an even stronger objection to Markowitz’s (1952) explanation, proving that no utility or probability weighting functions may explain the observed peanuts effect, unless the separability assumption is loosened.

#### Value-dependent dimension importance

What is difficult to explain in the peanuts effect is not just the existence of an interaction between value and probability (which issue may be readily fixed by loosening the assumption of independence), but the particular direction of the interaction. Parallel to decision under risk, there is an interaction effect between value and delay in temporal discounting, where humans discount future values less for higher values (Prelec & Loewenstein, 1991). This magnitude effect is explained by Prelec and Loewenstein’s (1991) decision theory, which assumes that higher values on one dimension increase the relative importance of the dimension in decision making. It may also be understood from the resource-rational perspective (Gershman & Bhui, 2020), where discounting is assumed to be consequence of noisy representation of future value, with higher values attracting more cognitive resources and thus associated with less discounting.

However, similar logic cannot explain the peanuts effect. Suppose for gambles of a higher stake, the value dimension is assigned a higher importance and accordingly the probability dimension a lower importance. Following this reasoning, the decision maker would be more attracted to the stake of the gamble and less sensitive to its risk. As a result, the theory predicts people would take more risks for larger gambles, which runs in the opposite direction of the peanuts effect.

#### Greater disappointment for larger gambles

That is why Prelec and Loewenstein’s (1991) decision theory could explain 9 out of 10 anomalies summarized in their paper but not the peanuts effect. This failure led Prelec and Loewenstein (1991) to resort to a distinctively different perspective to explain the peanuts effect, that people may maximize their own emotional experience instead of the expected utility, as formulated in the decision affect theory (Mellers et al., 1997). They reasoned that when the value of the gamble is higher, people would face greater disappointment for choosing the gamble but losing it. As a result, people may reverse their preference between gamble and sure reward when the value of the gamble increases. A similar idea was revisited in Weber and Chapman (2005).

Though intuitively appealing, this reasoning is missing the fact that the utility difference scales up with the disappointment when the value of the gamble becomes larger. For a smaller gamble where an individual prefers the gamble over the sure reward, the decision affect theory implies that the utility difference between the gamble and the sure reward outweighs the potential disappointment of choosing the gamble. When the gamble and the sure reward are proportionally amplified, the individual would switch their preference to the sure reward only if the disappointment function is an increasingly accelerating function of the utility difference. However, this does not receive empirical support from experiments measuring affective responses to gamble outcomes (Mellers et al., 1997). In fact, the empirical measurement of the affective response function even suggests a reverse effect.

#### Attention-dependent decision theories

There is increasing evidence from eye movement measurements that human decisions are influenced by the attention they allocate to available options. A natural question is whether the peanuts effect can be explained by attention-dependent decision theories such as the salience theory (Bordalo et al., 2012).

Fiedler and Glöckner (2012) found that the number of fixations to an outcome of a gamble increases with both its value and probability. He and Bhatia (2023) re-analyzed the eye movement data of Fiedler and Glöckner and further found that when the value of the recently fixated outcome is higher, participants are more likely to fixate on the winning probability of that outcome, indicating the cross-dimensional influence of value on the attention to probability. Consistent with these empirical findings, the salience theory (Bordalo et al., 2012) assumes that the decision maker’s attention is attracted to outcomes of higher value, and as the result, they would overweight probabilities accompanying higher values. Consequently, the salience theory would predict an effect in the reverse direction of the peanuts effect and thus cannot explain it.

### The two components of the peanuts effect

Though most previous studies (Binswanger, 1981; Estle et al., 2006; Kachelmeier & Shehata, 1992; Prelec & Loewenstein, 1991; Weber & Chapman, 2005) frame the peanuts effect as something about risk attitude, Fehr-Duda et al. (2010) find it to be more adequately described as a modulation of outcome value on the probability distortion function. A closer examination of the effect reveals two sub-effects (illustrated in Figure 1): (1) the sensitivity (slope) of subjective probability increases with value, and (2) the elevation (cross-over point) of subjective probability decreases with value.

**Figure 1.**
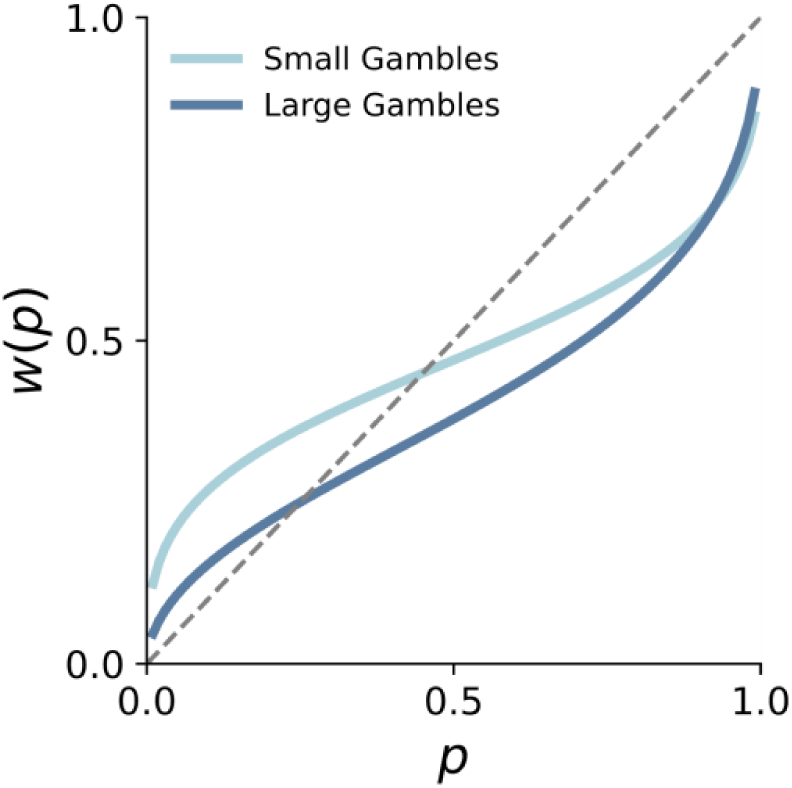
The Peanuts Effect on Probability Distortion. Illustration of the peanuts effect in decision under risk. Subjective probability (aka decision weight) *w*(*p*) is plotted as a function of objective probability *p*. Compared to that of small gambles (light blue curve), the subjective probability of large gambles (dark blue curve) has a steeper slope and a lower crossover point (i.e., where *w*(*p*) = *p*).

Though several resource-rational models have treated the distortion of probability (Bhui & Gershman, 2018; Zhang et al., 2020) or value (Heng et al., 2020; Polanía et al., 2019), the previous treatments are about probability or value alone and cannot explain why probability distortion may be influenced by the associated value. To explain such cross-modality influence, we extended previous work on resource-rational modeling and developed the ARRM framework.

## The assemblable resource-rational modules framework

One bottleneck of human cognitive processing is its limited resources which may be framed as information transmission through a constrained communication channel (Bhui et al., 2021; Dayan & Abbott, 2001; C. R. Sims, 2018; Wei & Stocker, 2015). Following previous work, we developed the Assemblable Resource-Rational Modules (ARRM) framework (see Figure 2A for an overview) to model resource-rational agents who use their limited communication channel adaptively to process the information needed for decision making.

**Figure 2.**
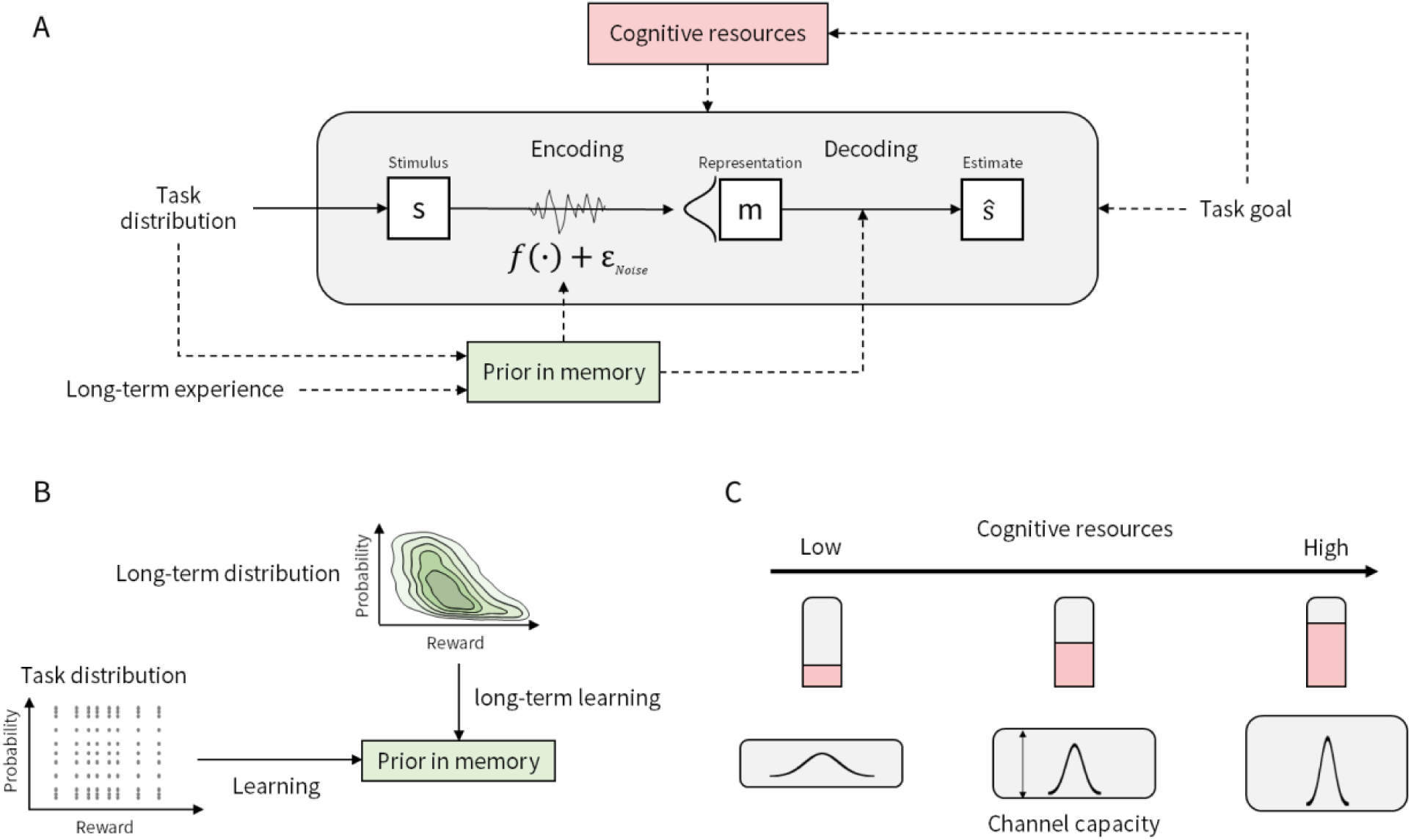
The ARRM framework of stimulus processing. (**A**) Schema of the framework, where a resource-rational agent’s encoding and decoding processes are constrained by limited cognitive resources and optimized under specific prior in memory and optimization goal (loss function). The allocation of cognitive resources may also be optimized. (**B**) Illustration of two possible origins of the prior information in memory: the stimulus distribution in the current task (task distribution) and the distribution from long-term life experience (long-term distribution). The prior distribution can either be a marginal distribution over the stimulus in question (e.g., the probability of an outcome), or a conjunctive distribution of multiple attributes (e.g., the conjunctive distribution of value and probability), as implied by the saying “the higher the risk, the greater the return” (illustrated in the plot for long-term distribution). (**C**) Illustration of channels with different levels of cognitive resources. Higher cognitive resources lead to higher representational precision (i.e., lower noise).

Without loss of generality, the processing of a specific stimulus can be decomposed into an encoding process (from stimulus s to a noisy internal representation *m*) and a decoding process (from representation *m* to a subjective estimate *ŝ*). The encoding and decoding schemes chosen by a resource-rational agent may depend on the agent’s cognitive resources, prior beliefs about the environment, and optimization goal (loss function). These “modules” together constitute the ARRM framework. The framework has multiple variants for each module, and as the “assemblable” in its name suggests, allows the variants from different modules to be combined in an unconventional way to form new resource-rational models. Below we specify the possible variants for each module, focusing on how the model space may be systematically expanded to cover a broader range of decision phenomena.

### Cognitive resource module: three allocation problems

Resource-rational models of decision making share the assumption that the brain optimizes information transmission through a constrained communication channel, but differ in their assumptions about resource constraints or optimization goals. These models descend from two theoretical lines: efficient coding (Attneave, 1953; Barlow, 1961; Simoncelli & Olshausen, 2001; Wei & Stocker, 2015) and the rate-distortion theory (C. J. Bates et al., 2019; Gershman & Bhui, 2020; MacKay, 2003; C. R. Sims, 2016). The resource assumptions in these two lines of models are analogous to the charges in two types of mobile data plans: Efficient-coding models resemble plans with a fixed monthly fee, where each bit of transmitted information counts as a gain; in contrast, rate-distortion models are like plans charged by data usage, where each bit of information adds to the cost. Regardless of how cognitive resources are defined, the higher the resource used to encode a stimulus, the more precise the representation (Figure 2C).

Resource-rational agents face three types of allocation problems. The first is the classic coding problem of how to allocate cognitive resources across different possible values of the to-be-encoded stimulus. This is also the problem most of previous efficient coding models (Bhui & Gershman, 2018; Polanía et al., 2019; Wei & Stocker, 2015) focus on, that is, to adjust the encoding scheme according to the prior distribution (or information source) of the stimulus to fulfill specific goals.

The second problem considers a more dynamic view: How to allocate cognitive resources available across time? In 2003, C. A. Sims proposed the hypothesis of *rational inattention* that people should invest more of their limited cognitive resources in choices that are more important (C. A. Sims, 2003), which has been used to model a variety of decision phenomena (Gershman & Bhui, 2020; Grujic et al., 2022; Mikhael et al., 2021; C. A. Sims, 2003). The rational inattention hypothesis is also consistent with qualitative observations that people tend to take more cognitive effort when the task incentives increase (Kleinsorge & Rinkenauer, 2012; Otto & Vassena, 2021). One difference between the first (classic-coding) and the second (dynamic-view) resource-allocation problems is that the former is only about the to-be-encoded stimulus itself, whereas the latter may allow influences from other stimulus dimensions, such as allocating more resources to a probability stimulus when it is in a gamble associated with a more valuable outcome (Figure 2C), potentially relevant to cross-modality effects like the peanuts effect.

The third problem addresses resource allocation among different dimensions in multi-dimensional decision-making scenarios, such as allocating resources between value and probability in a gamble. Rational agents should allocate more resources to dimensions that are more uncertain (Dewan, 2020) or contribute more to the task goal (C. J. Bates et al., 2019; C. J. Bates & Jacobs, 2021). Though we will not cover such cross-dimension resource allocation in our modeling of the peanuts effect, it is an important part of real-world decisions that can be integrated into resource-rational modeling.

### Priors in memory module: two sources of distributions

A resource-rational agent must also consider the stimulus distribution of the environment. More precisely, it is the agent’s subjective beliefs about the distribution instead of the objective distribution that defines its optimization problem. To emphasize that these prior beliefs are maintained in memory and themselves resource-constrained, we call them *priors in memory*.

These priors can be based on the stimulus distribution in the current task environment (task distribution) or the long-term environmental statistics (long-term distribution). While using priors learned from the task distribution seems self-evident for maximizing task performance (Heng et al., 2020; Jazayeri & Shadlen, 2010), using priors based on the long-term distribution is also justified by accounting for a wide range of human behaviors in judgment and decision making (Girshick et al., 2011; Griffiths & Tenenbaum, 2006; Stewart et al., 2015; Wei & Stocker, 2015), especially when learning the stimulus distribution separately for each task is costly or implausible due to limited samples (Binz et al., 2022; Dasgupta et al., 2020).

Using the conjunctive distribution of reward and probability in gambles to illustrate these two sources of priors (Figure 2B), we emphasize two facts. First, these priors are not necessarily only about one stimulus dimension, as widely assumed in previous resource-rational models, but may reflect the covariation of multiple dimensions. Second, even when reward and probability are independent in the task distribution, the long-term distribution from life experience may exhibit a negative correlation between reward and probability (Pleskac & Hertwig, 2014), echoing “the higher the risk, the greater the return”. Such structural long-term prior has the potential to explain the interdependence between value and probability in the peanuts effect.

### Possible variants of other modules

Having discussed the modules of cognitive resources and priors in memory, whose extensions are essential to explaining the peanuts effects, we now briefly summarize the possible variants of other modules in the ARRM framework.

*Encoding schemes*, which map stimuli to internal representations, can be classified into three levels of adaptiveness based on their flexibility in adapting to cognitive resources and priors in memory: fixed, boundedly-adaptive, and fully-adaptive. Fixed encoding schemes, assumed in traditional Bayesian observer models (Petzschner et al., 2015), do not change with cognitive resources or priors. Boundedly-adaptive encoding schemes, such as the Bounded Log-Odds (BLO) model (Ren et al., 2021; Zhang et al., 2020), may adapt to cognitive resources or priors but in a constrained way, with fixed functional forms and adaptive parameters. Fully-adaptive encoding schemes, used in many efficient coding models and rate-distortion models (Bhui & Gershman, 2018; Heng et al., 2020; Wei & Stocker, 2015), allow for flexible changes in the encoding mapping and precision based on the prior distribution and available cognitive resources. The adaptation of encoding schemes to the environment may occur on multiple time scales, with slow adaptation to long-term environmental statistics shaping the functional form and fast adjustments within that form. While the speed of adaptation to the task distribution in general is still largely unknown, there is stronger evidence for trial-by-trial changes in encoding precision (C. J. Bates & Jacobs, 2021; Gershman & Bhui, 2020).

*Decoding schemes*, while omitted in some efficient coding models (Bhui & Gershman, 2018; Heng et al., 2020; Stewart et al., 2006), are necessary for cases where a stimulus needs to be explicitly estimated or integrated with stimuli on other dimensions (Polanía et al., 2019; Wei & Stocker, 2015; Zhang et al., 2020), such as the represented probability is to be integrated with value. For a rational agent, a statistically principled decoding method is Bayesian inference (Petzschner et al., 2015; Wei & Stocker, 2015), which combines the noisy representation with prior information to calculate the posterior distribution of the stimulus. Alternatively, in rate-distortion models, the mapping from the stimulus to its estimate can be directly obtained based on the rate-distortion theory, without specifying the details of encoding and decoding. Even so, a Bayesian decoding process is implicitly assumed, because it constitutes the only optimal solution to the optimization problem.

The optimization goal (or loss function) is indispensable for defining any resource-rational agent. The loss function can be a function of the deviation between stimulus and its subjective estimate, or refers to the loss that is explicitly defined by the task (e.g., error rate or monetary loss). In previous efficient coding models, the goal is often to maximize the mutual information between stimulus and representation (Bhui & Gershman, 2018; Wei & Stocker, 2015). Using the explicit task goal to define loss function is common in previous rate-distortion models (C. J. Bates & Jacobs, 2021) but less popular in previous efficient coding models (see Heng et al., 2020 for an exception), though it is applicable to both lines of resource-rational models.

Solving the exact optimization problem can be difficult or even intractable for the brain, especially in complex environments and on fast time scales. As a result, humans may adapt to the environment in a more bounded way than perfect resource-rational agents. The ARRM framework can model this boundedness in several ways. First, the encoding scheme may be only partially adaptive to the environment, as assumed in the bounded log-odds model (Zhang et al., 2020), and approximate implementations of resource rationality, such as the Decision by Sampling model (Stewart et al., 2006), may better capture human behavior than exact resource-rational agents (Heng et al., 2020). Second, the encoding scheme typically adapts to the environment over the long run (Stewart et al., 2006; Wei & Stocker, 2015) rather than on shorter time scales of minutes or hours (Alempaki et al., 2019; Bujold et al., 2021; Frydman & Jin, 2021; Ren et al., 2018), while the adaptation of the decoding scheme may be much faster, as suggested by the Bayesian literature (Maloney & Zhang, 2010; Petzschner et al., 2015). What is explored in the ARRM framework above are theoretical possibilities, while how humans may behave like resource-rational agents is an empirical question. Next, we apply the framework to decision under risk to explain the peanuts effect and examine which set of theoretical possibilities best fits human behaviors.

## ARRM MODELS FOR EXPLAINING THE PEANUTS EFFECT

In the following, we apply the ARRM framework to modeling human decision under risk. By combining the rational inattention hypothesis of cognitive resource allocation (Gershman & Bhui, 2020; Grujic et al., 2022; C. A. Sims, 2003) with a hypothesis of value-probability covariation in prior beliefs (Pleskac & Hertwig, 2014), we are able to provide a full explanation for the peanuts effect in probability distortion.

We use ℊ: (*x*_1_, *p*; *x*_2_) to denote a monetary gamble with two outcomes, that is, receiving *x*_1_ with probability *p*, or *x*_2_ otherwise, where |x_1_| > |x_2_|. A sure payoff (gain or loss) is a special case of this two-outcome gamble with *p* = 1. Following CPT (Tversky & Kahneman, 1992), we assume that the expected utility of gamble ℊ can be computed as^2^

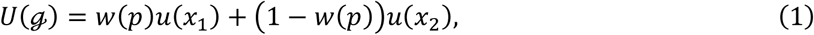

where *u*(·) denotes the utility function and *w*(·) denotes the probability weighting (distortion) function. What differs our following modeling from standard applications of CPT is that we replace the probability weighting function *w*(·) of *p* with the subjective estimate *p̂*, which is decoded from a noisy representation of *p* and may vary from gamble to gamble as the consequence of resource-rational encoding and decoding.

### The Rational Inattention (RI) hypothesis

As shown in Figure 2C, the resource-allocation problem can be understood as a tradeoff between task performance and cognitive effort: A higher encoding precision would lead to a more accurate estimation and thus better task performance, but at a cost of higher demand for cognitive resources. To balance task performance and cognitive effort, a resource-rational agent should allocate cognitive resources among different stimuli according to their potential rewards. In behavioral economics, this is known as the rational inattention (RI) hypothesis (Grujic et al., 2022; Matějka & McKay, 2015; Mikhael et al., 2021; C. A. Sims, 2003). It addresses the problem of how to allocate cognitive resources over time to maximize expected reward (i.e., the dynamic allocation problem): stimuli associated with a larger reward receive more attention, while those that are worthless are ignored (Kool & Botvinick, 2018; Matějka & McKay, 2015).

When applied to the representation of probability, the RI hypothesis implies that the cognitive resource increases with the potential gain of the gamble. For gamble (*x*_1_, *p*; *x*_2_), the resource allocated to the probability *p* should increase with its associated potential gain (or saved loss) |x_1_ − x_2_|:

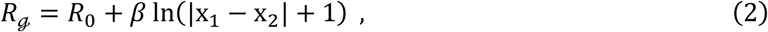

where *R*_0_ ≥ 0 is a parameter for the baseline resource (i.e., when x_1_ − x_2_ = 0), and *β* ≥ 0 is a slope parameter controlling how steep the allocated resource changes with value. This assumption of linear relationship is an approximation to the optimal resource allocation (see Supplemental Text S3 and Figure S3 for more details). Similar linear-log forms of rational inattention assumptions have been used in previous studies (Gershman & Bhui, 2020; Mikhael et al., 2021).

### The Structural Prior (SP) hypothesis

According to the module of priors in memory of the ARRM framework, one source for prior beliefs is the long-term experience in everyday life (figure 2B). Such everyday statistics have been used by previous resource-rational models to explain human distortion of value, probability or time (Bhui & Gershman, 2018; Stewart et al., 2006, 2015), but the priors used in these practices are limited to one single dimension of stimuli.

However, as the saying “the higher the risk, the greater the return” suggests, a structural relationship of reward and probability—the two dimensions of a gamble—widely exists in everyday life, ranging from financial investment to the submission of academic papers (Pleskac & Hertwig, 2014). That is, when the potential value is large, the probability of success tends to be small. Indeed people may use value as a contextual cue to infer the associated probability (Leuker et al., 2018, 2019). In the structural prior (SP) hypothesis, we hypothesize that people have prior beliefs about the probabilistic interdependence between value and probability.

Following previous studies (Khaw et al., 2021), we model the agent’s prior belief for *p* as a Gaussian distribution on the log-odds scale:

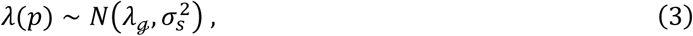

where 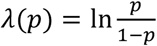 is the log-odds of *p*. The *λ*_ℊ_ and 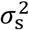 respectively denote the mean and variance of the Gaussian prior, with the subscript ℊ of *λ*_ℊ_ to emphasize its dependence on the gamble in question. When transformed back to the probability scale, the prior would be *U*-shaped for large variance and inverted-*U*-shaped for small variance, resembling the beta distribution used to model the real-world statistics of probability (Stewart et al., 2006; Zhu et al., 2020).

The SP hypothesis for gamble ℊ: (*x*_1_, *p*; *x*_2_), can then be formulated as

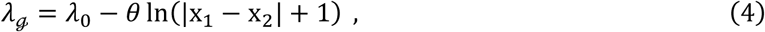

where *λ*_0_ is a baseline parameter, |x_1_ − x_2_| (as before) is the potential gain (or saved loss) of the gamble, and *θ* is a slope parameter controlling the direction and extent that probability is expected to change with value, with *θ* > 0 indicating an assumed negative correlation between probability and value, *θ* = 0 for no correlation, and *θ* < 0 for positive correlation.

### Probability estimation models

To implement the resource rational framework for probability estimation, we adopt three previous resource-rational decision models as the “base” models and combine them with the RI and SP hypotheses (“modifiers”) to produce a set of 10 different models. Below we briefly review each of the base models and how they may be enhanced by the RI and SP hypotheses (see Tables 1 and 2 for a summary of model notations).

**Table 1.**
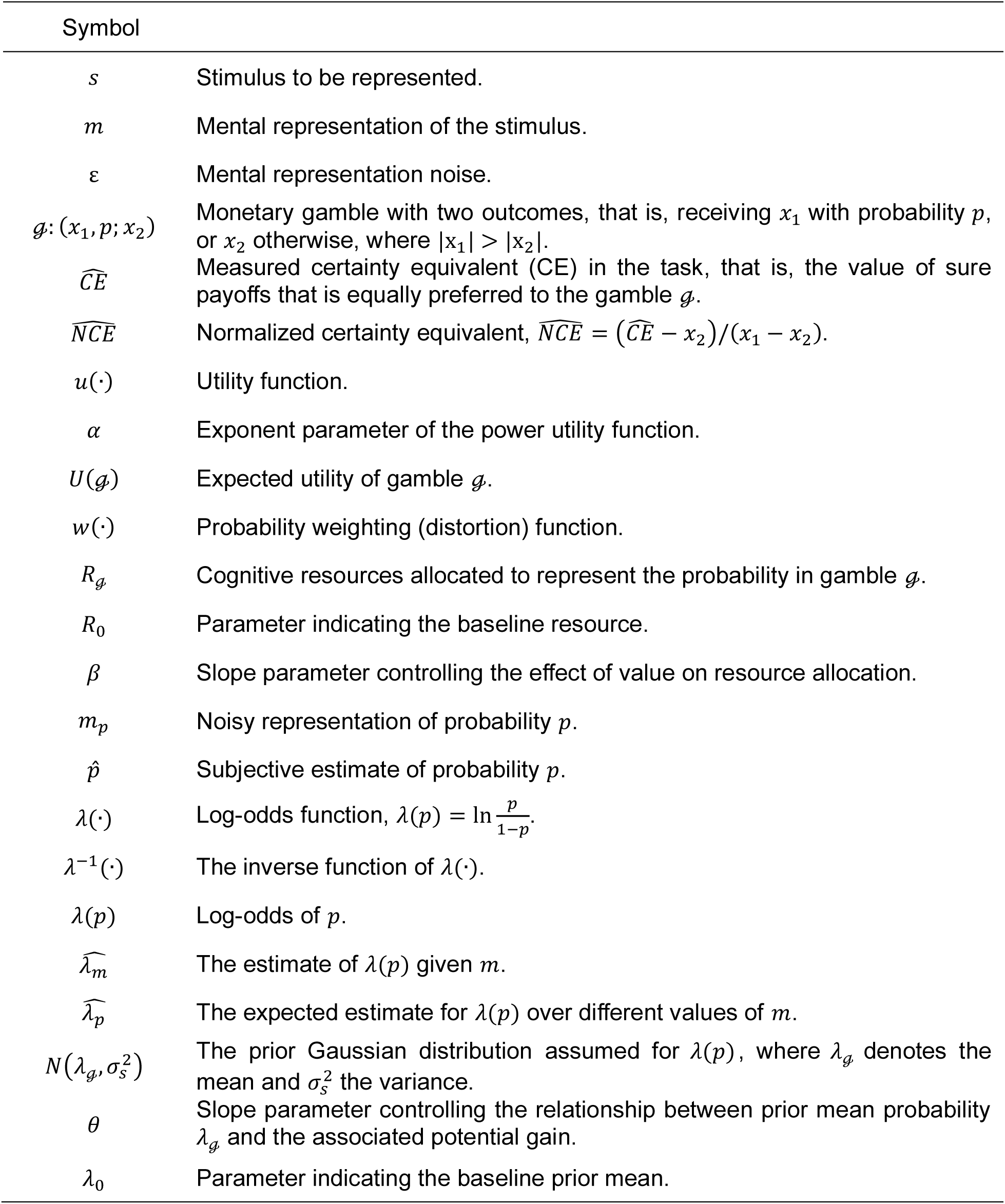
General Notations.

**Table 2.**
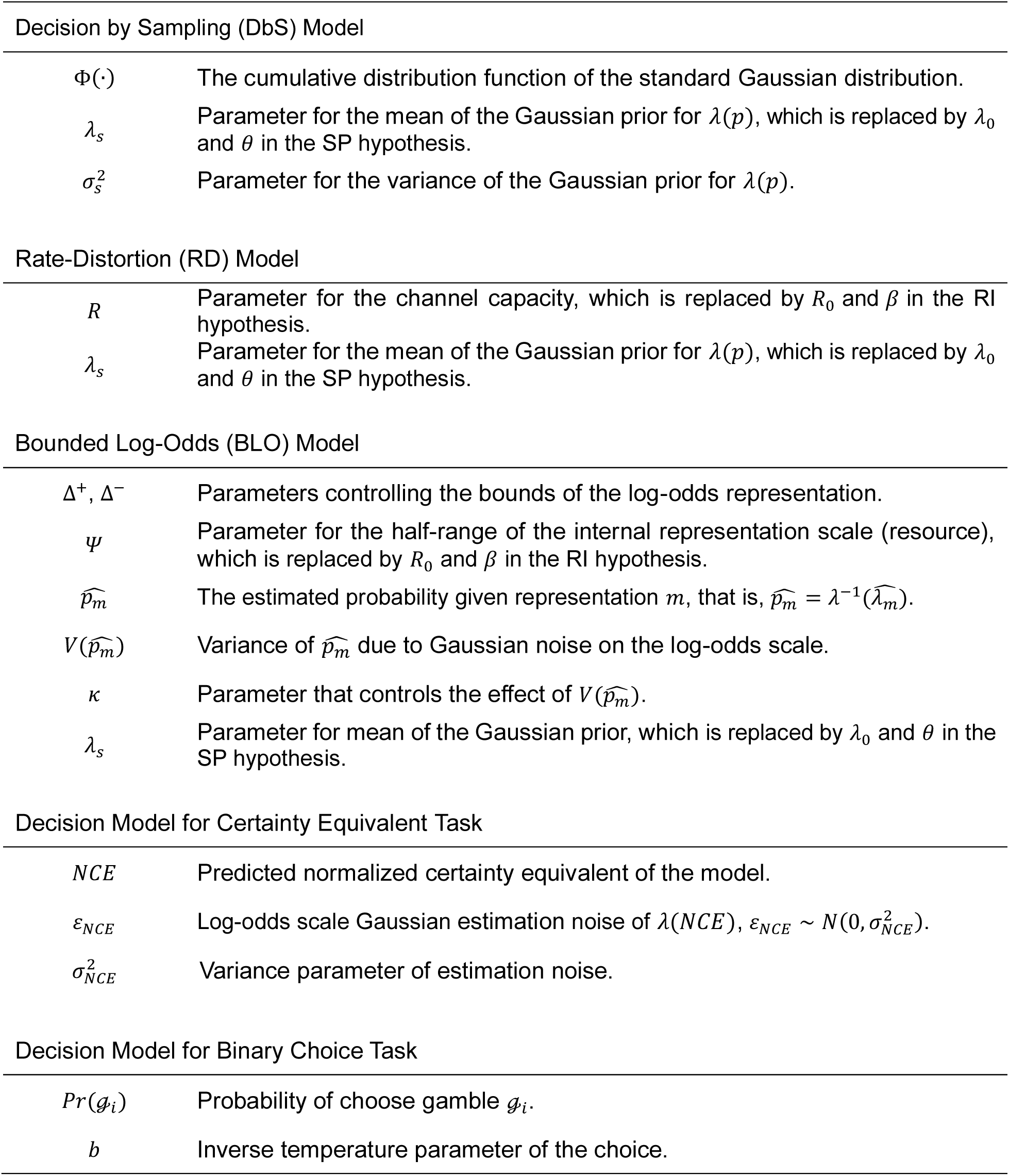
Model Notations.

#### Decision by Sampling (DbS) model

The DbS model (Bhui & Gershman, 2018; Stewart et al., 2006) belongs to the theoretical line of efficient coding, assuming fully-adaptive encoding scheme, no decoding scheme and the optimization goal of maximizing mutual information. As its name suggests, it starts with sampling from memory. It assumes that the agent compares the stimulus with each sample and calculates the proportion of times the former is larger than the latter as the subjective estimation of the stimulus. When sample size is sufficiently large, this subjective estimation approximates the cumulative distribution function of the prior distribution of the samples (Bhui et al., 2021; Bhui & Gershman, 2018; Stewart et al., 2015). Given prior λ(p) ∼ *N*(*λ*_*s*_, 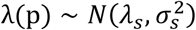) (Eq. 3, with *λ*_ℊ_ set to a constant *λ*_*s*_), the subjective estimation of probability p can be written as:

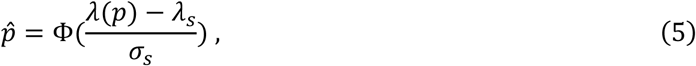

where Φ(·) denotes the cumulative distribution function of the standard Gaussian distribution.

There is no obvious way that the DbS model can be enhanced by the RI hypothesis, because its subjective estimation (Eq. 5) is not influenced by the available cognitive resource (in fact, the model assumes a sufficiently large sample size to approximate the cumulative probability function). Therefore, we only consider the SP hypothesis for DbS, which is simply to replace the *λ*_*s*_ in Eq. 5 with the *λ*_ℊ_ in Eq. 4. In the special case of *θ* = 0, it reduces to the original DbS without the SP hypothesis. The original DbS model has two free parameters: *λ*_*s*_, and *σ*_*s*_. The DbS+SP model replaces the parameter *λ*_*s*_ with two parameters: *λ*_0_ and *θ*. Unless otherwise specified, our count of model parameters does not include the parameters of the decision model (introduced later), which are shared across all 10 models.

#### Rate-Distortion (RD) model

The RD model assumes a constraint on the rate of transmitted information and optimizes encoding and decoding schemes under this constraint to minimize the expected loss incurred by distorted subjective estimation of the stimulus. It has a fully-adaptive encoding scheme and a Bayesian decoding scheme, whose optimization goal may be defined by the task.

Similar to previous RD models (C. J. Bates et al., 2019; Gershman & Bhui, 2020), we assume a Gaussian information source and a squared-error loss function, for which the optimal encoding scheme is a linear mapping with homogeneous Gaussian noise (Berger & Gibson, 1998). We further assume that the encoding of probability is on the log-odd scale (Khaw et al., 2021; Zhang et al., 2020; Zhang & Maloney, 2012), so the representation can be written as

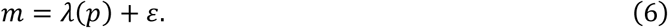

Here *λ*(*p*) is the log-odds of probability *p*, and 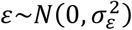 is a noise term with mean 0 and variance 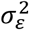.

The optimal decoding scheme is Bayesian decoding. Given the Gaussian prior 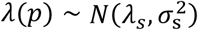 and representation *m*, the posterior estimate of *λ*(*p*) follows a Gaussian distribution:

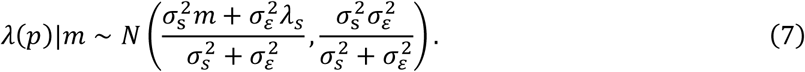

Given *m*, the optimal estimate for *λ*(*p*) that minimizes squared error on the log-odds scale is the posterior mean,

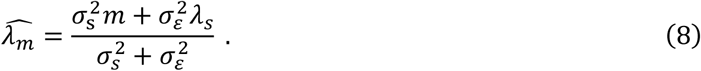

Marginalizing out *m*, we have the expected estimate for *λ*(*p*):

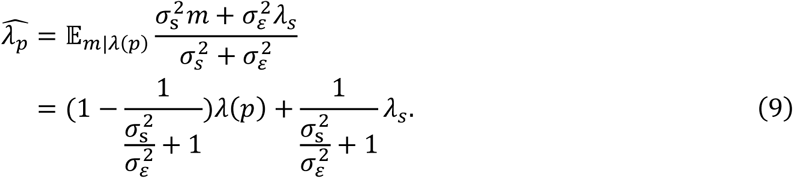

According to information theory, for a Gaussian source (variance 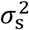) transmitted through a Gaussian channel (variance 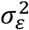), the rate of information flow, *R*, equals the channel capacity, which satisfies

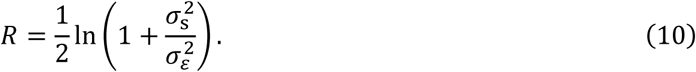

Substituting Eq. 10 into Eq. 9, we have

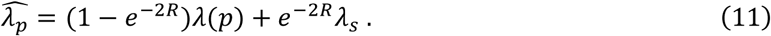

This form is consistent with Linear in Log-Odds—a two-parameter descriptive model of probability distortion widely used in previous studies (Gonzalez & Wu, 1999; Zhang & Maloney, 2012)—and provides a cognitive explanation for its two parameters. The cognitive resource *R* controls the sensitivity parameter (slope) of the distortion curve, with higher resource associated with higher sensitivity (steeper slope); the prior *λ*_*s*_ controls the elevation parameter (intercept) of the curve.

The combinations of the RD model with the RI and SP hypotheses are straightforward, which respectively replace the *R* and *λ*_*s*_ in Eq. 11 with the *R*_ℊ_ in Eq. 2 and the *λ*_ℊ_ in Eq. 4. The original RD model has two free parameters: *R* and *λ*_*s*_. The RD+RI and RD+RI+SP models replace the parameter *R* with two parameters: *R*_0_ and *β*. The RD+SP and RD+RI+SP models replace the parameter *λ*_*s*_ with two parameters: *λ*_0_ and *θ*.

#### Bounded Log-Odds (BLO) model

Similar to DbS, the BLO model (Ren et al., 2021; Zhang et al., 2020) is an efficient coding model and assumes mutual information maximization as the optimization goal. But different from DbS, it has a boundedly-adaptive encoding scheme that uses fixed functional forms and adaptive parameters to approximate optimal encoding solutions, and an approximately Bayesian decoding scheme.

The model assumes an agent who encodes probability in log-odds on an internal scale with finite length and homogeneous Gaussian noise. Define a bounded transformation of log-odds

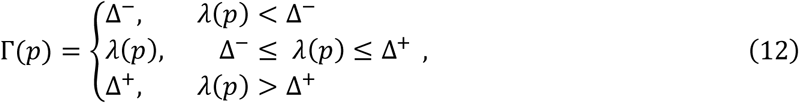

where *λ*(*p*) (as before) is the log-odds of *p*, and Δ^−^ and Δ^+^ are the bounds parameters specifying the encoded stimulus range. A mapping of the transformed Γ(*p*) to the internal representation scale [−*Ψ*, *Ψ*] is:

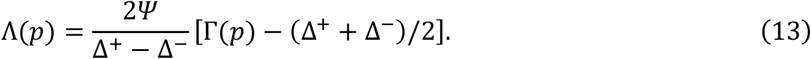

The noisy internal representation is

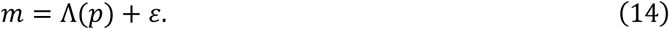

where 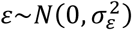 is a noise term with mean 0 and variance 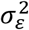.

A subjective log-odds 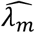 can be decoded from the representation *m* and further transformed back to the probability scale to obtain 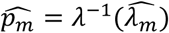. The decoding goal of the BLO model is not to minimize the squared error for 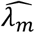 on the log-odds scale, but to minimize the squared error for 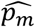 on the probability scale, whose Bayesian inference has no known analytical form. The BLO model adopts the following form to compensate for the representational noise:

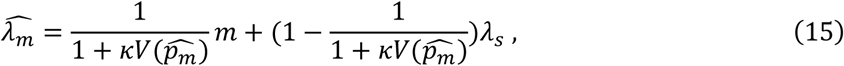

where *λ*_*s*_, as before, is the mean of the Gaussian prior, ***κ*** is a free parameter, 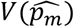 is the variance of 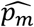 due to Gaussian noise on the log-odds scale, whose approximation has the form:

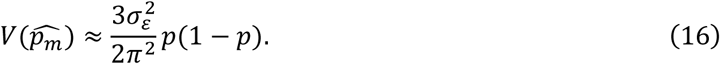

The agent is assumed to have a separate estimate of this variance without knowing the exact value of *p*. The parameter *σ*_*ɛ*_ is redundant with ***κ*** and thus omitted in model fitting.

Similar to the proof for Eq. 9, we can write the expected estimate of *λ*_*m*_ given *p* as

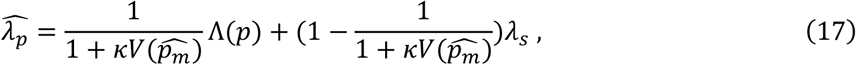

To apply the RI hypothesis to BLO, we replace *Ψ* with the *R*_ℊ_ in Eq. 2. Similar to the enhancement of the DbS and RD models, the SP hypothesis is applied to BLO through replacing the *λ*_*s*_ in Eq. 17 with the *λ*_ℊ_ in Eq. 4.

### Decision model

All the probability estimation models above are combined with the same decision model to produce responses in a specific task. We assume a power utility function

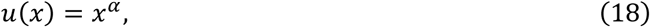

where *⍺* > 0 is a free parameter, which may be different for gain and loss. Substituting it into Eq. 1 and substituting the probability weighting function *w*(*p*) in Eq. 1 with the subjective estimate *p̂*, the expected utility of gamble ℊ can be written as

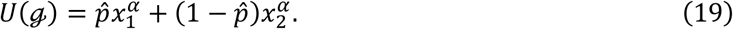

where *p̂* follows Eq. 5 for the DbS model and computed as

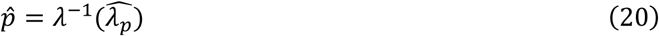

for the RD and BLO models.

Two typical tasks of decision under risk, certainty equivalent task and binary choice task, are modeled as the following.

#### Certainty Equivalent Task

On each trial, participants choose between a gamble and an array of sure payoffs of varying values so that their certainty equivalent (CE)—the value of sure payoffs that is equally preferred—to each gamble can be estimated for individual participants. The decision model above predicts

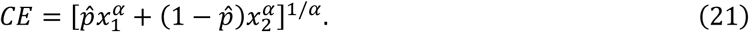

We define normalized certainty equivalent

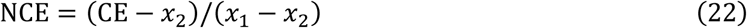

as a measure of subjective probability and compute a log-odds transformation of the measure for both participants’ and model-predicted NCEs. Participants’ 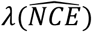 is modeled as model prediction plus a Gaussian noise:

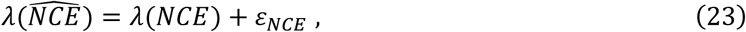

where 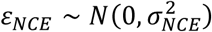 and *σ*_*NCE*_ is a free parameter.

#### Binary Choice Task

On each trial, participants choose between two gambles, or between one gamble and one sure payoff. We model the probability of choosing gamble ℊ_*i*_ as a SoftMax function determined by its utility difference with the other gamble:

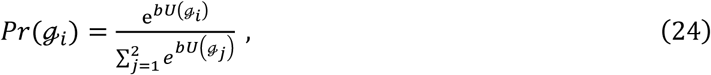

where *b* is an inverse temperature parameter that controls the stochasticity of the choice. Participants’ binary choice on each trial is modeled as a Bernoulli random variable based on *Pr*(ℊ_*i*_).

## Study 1: modeling the peanuts effect in the ARRM framework

To see how well the resource-rational models we constructed in the ARRM framework can explain human decision behaviors, we first need a model-free quantification of the phenomenon in question. For the peanuts effect, which reflects the influence of outcome value on probability distortion (Fehr-Duda et al., 2010), the ideal experimental design shall allow us to measure the probability distortion curve separately for different values. Fortunately, two previously published datasets satisfy our requirement, one from Gonzalez and Wu (1999) and the other from Zhang et al. (2020), which used an almost identical design of 165 different two-outcome gambles from a factorial combination of 11 different probabilities and 15 different sets of values (see Methods in Appendix). For each of the 10 or 75 participants, the authors of the studies measured the certainty equivalent (CE) for each gamble, that is, the sure reward that is perceived as equally preferable as the gamble. Based on these certainty equivalents, we elicited the probability distortion curve for each set of values and quantified the two aspects of the peanuts effect—how the sensitivity and elevation parameters of probability distortion change with value.

### The peanuts effect in the GW99 and ZRM20 datasets

Non-parametric estimation of probability distortions had been performed for the GW99 (Gonzalez & Wu, 1999) and ZRM20 (JDA and JDB) datasets (Zhang et al., 2020) in their original studies. Such estimation, however, had collapsed data across gambles with different outcome values.

The normalized certainty equivalent, NCE = (CE − *x*_2_)/(*x*_1_ − *x*_2_), was used as a measure of subjective probability, which could be computed for each gamble based on the estimated CE for the participant (denoted 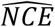 as estimation). Though 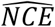 is not strictly equal to subjective probability unless the utility function is linear, it served as a useful measure to visualize the influence of outcome value on probability distortion. Figure 3 plots the mean 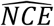 across participants for each dataset (GW99, n=10) or sub-dataset (JDA, n=51; JDB, n=24), separately for gambles with different *x*_1_ (Only gambles with *x*_2_ = 0 is plotted for visualization purposes).

**Figure 3.**
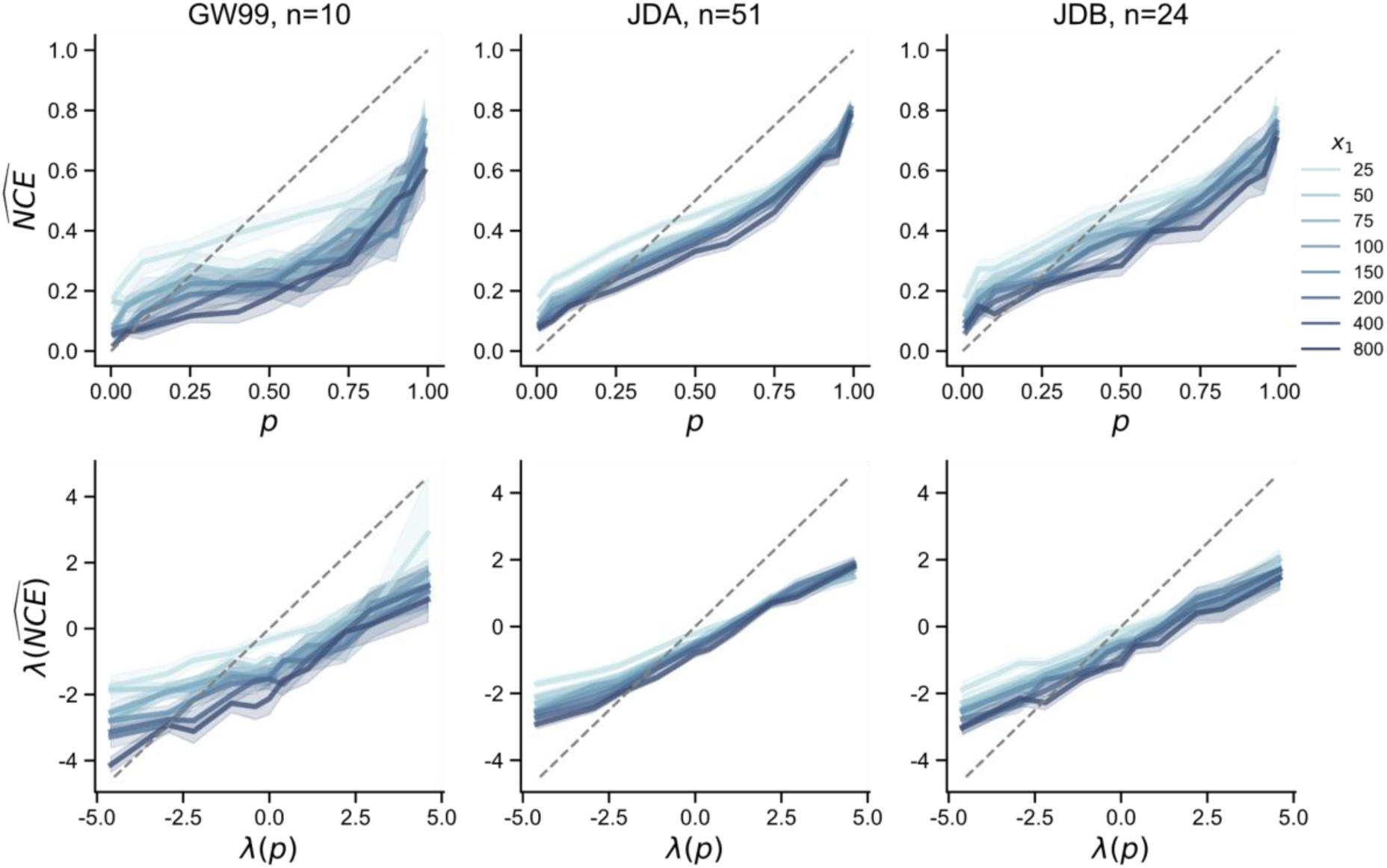
The peanuts effect in the GW99 and ZRM20 (JDA and JDB) datasets. For each dataset or sub-dataset (one column), participants’ normalized certainty equivalent (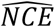, a measure of subjective probability) is plotted against objective probability (*p*), separately visualized for gambles (*x*_1_, *p*; 0) with different *x*_1_ (color-coded in different darkness). The upper and lower rows are respectively for probability and log-odds plots. Shadings denote standard error. Both sub-effects of the peanuts effect are visible in all the datasets: the larger the gamble (darker curves), (1) the greater the sensitivity of subjective probability, and (2) the lower the elevation of subjective probability.

The observations in all these datasets confirmed the peanuts effect and its two sub-effects, whose descriptions had been scattered in different previous studies (Fehr-Duda et al., 2010; Green et al., 1999; Prelec & Loewenstein, 1991). Other than exhibiting the typical inverted *S*-shape (linear in the log-odds scale) as in the original analyses of the datasets, the probability distortion curve had (1) *greater sensitivity* (i.e., steeper slope) and (2) *lower elevation* (i.e., smaller crossover point) for larger gambles (darker curves in Figure 3). The sensitivity effect and elevation effect were supported by linear mixed-effect model (LMM) analysis for participants’ 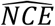 (Eq. 11 in Methods of Appendix). When the potential gain (*D* = |*x*_1_ − *x*_2_|) was greater, participants were more sensitive to the differences between different probabilities (*D* by probability interaction, GW99: *b* = 0.084, 95% CI [0.055, 0.114], *P* < 0.001; JDA: *b* = 0.067, 95% CI [0.060, 0.074], *P* < 0.001; JDB: *b* = 0.050, 95% CI [0.019, 0.081], *P* = 0.002), and had lower 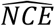 (main effect of *D*, GW99: *b* = −0.211, 95% CI [−0.270, −0.153], *P* < 0.001; JDA: *b* = −0.090, 95% CI [−0.114, −0.066], *P* < 0.001; JDB: *b* = −0.093, 95% CI [−0.131, −0.055], *P* < 0.001).

### Exchangeability test: rejecting the separability assumption of value and probability

If participants’ choices had followed the prospect theory with power utility function *u*(*x*) = *x*^*⍺*^ and any probability weighting function *w*(*p*), for gamble (*x*_1_, *p*; *x*_2_) with *x*_2_ = 0, we would have *NCE* = *w*(*p*)^*⍺*^. If so, *NCE* would simply be a function of *p* and have nothing to do with outcome value. However, as shown in the sensitivity and elevation effects above, *NCE* changed systematically with outcome value. In other words, the peanuts effect violates the separability assumption widely accepted in previous decision models—utility is only determined by value and subjective probability only by probability. This has motivated us to develop resource-rational models to capture the cross-dimensional influence of outcome value on probability distortion.

But one may wonder whether some non-power utility function, such as the four-fold function proposed by Markowitz (1952), could explain the peanuts effect without modifying the separability assumption. To exclude this possibility, we developed a model-free test for separability, termed the *exchangeability test*, whose conclusion can apply to any forms of utility and probability weighting functions.

The basic idea of the test is to navigate in a space of gambles through two different paths, analogous to first going west then north vs. first north then west in the physical space. If the separability between value and probability held, the two paths would lead to the same destination. In particular, the test (illustrated in Figure 4A) is based on the CE measured in the GW99 and ZRM20 datasets for different gambles (Figure 3). For gamble (*x*_1_, *p*; 0), the separability assumption implies

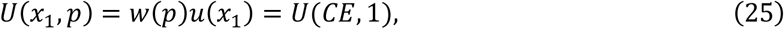

**Figure 4.**
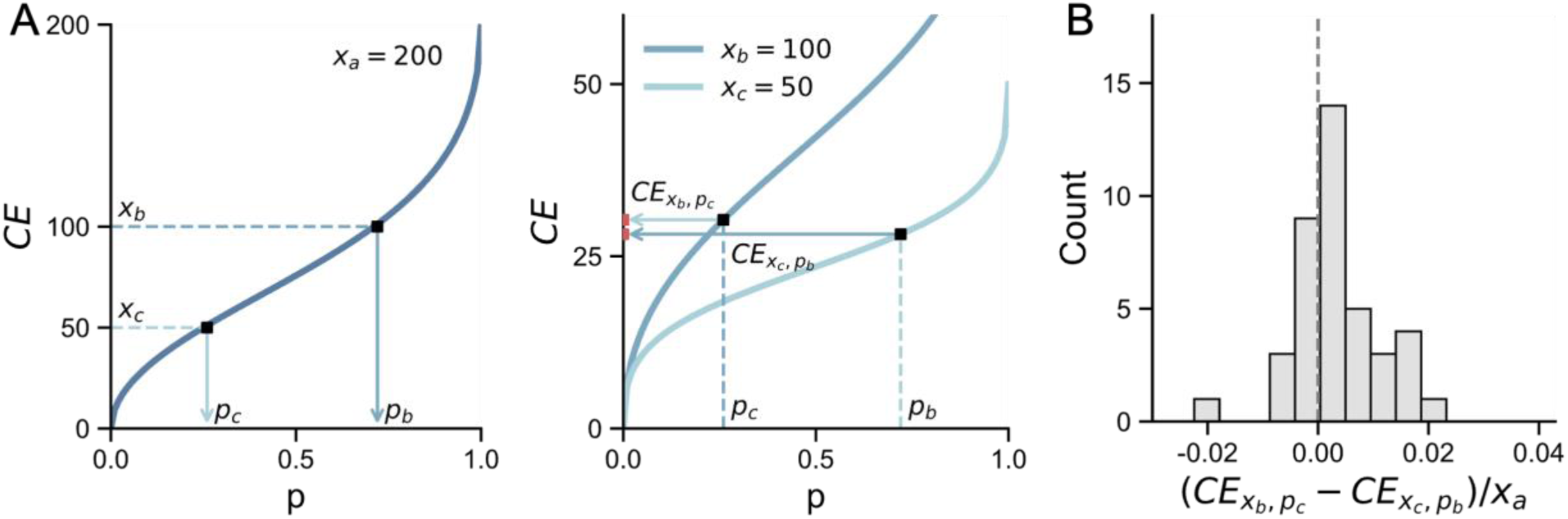
The exchangeability test in the GW99 and ZRM20 datasets. (**A**) Illustration of the rational of the exchangeability test. The test was based on the 8 certainty equivalents (CE) curves estimated from the datasets, separately for gambles with *x*_1_ = 25, 50, 75, 100, 150, 200, 400, 800. From these 8 different values of *x*_1_, we drew 40 combinations of *x*_*a*_, *x*_*b*_, *x*_*c*_ that satisfy (*x*_*a*_, *p*_*b*_; 0)∼*x*_*b*_, (*x*_*a*_, *p*_*c*_; 0)∼*x*_*c*_, and 0.01 ≤ *p*_*b*_, *p*_*c*_ ≤ 0.99, where ∼ denotes “equally preferred” (**left panel**). We then tested whether the equivalence pair predicted by the separability of value and probability—(*x*_*c*_, *p*_*b*_; 0) and (*x*_*b*_, *p*_*c*_; 0)—were equivalent in CE (**right panel**). (**B**) Results of the exchangeability test: the histogram of the 40 equivalence pairs’ CE differences. The mean 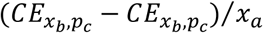 of the 40 equivalence pairs were significantly greater than 0.

where *U*(*x*_1_, *p*) is the expected utility of the gamble, and *U*(*CE*, 1) is the utility of the CE. For any *x*_*a*_ > *x*_*b*_ > *x*_*c*_, we could find probabilities *p*_*b*_ and *p*_*c*_ so that the CE of gamble (*x*_*a*_, *p*_*b*_; 0) is *x*_*b*_ and the CE of gamble (*x*_*a*_, *p*_*c*_; 0) is *x*_*c*_. Using these two probabilities we can construct two new gambles (*x*_*c*_, *p*_*b*_; 0) and (*x*_*b*_, *p*_*c*_; 0). It is easy to prove that these two gambles have the same CE (by applying Eq. 25). Significant differences between the two gambles would allow us to reject separability (see Methods in Appendix for more details).

The GW99 and ZRM20 datasets allowed us to test 40 pairs of equivalence, across which the measured CE for (*x*_*c*_, *p*_*b*_; 0) and (*x*_*b*_, *p*_*c*_; 0) were significantly different from each other (paired *t*-test, *t*(39) = 2.79, *p*=0.008), thus rejecting the separability assumption (Figure 4B). That is, no utility or probability weighting functions would explain the observed peanuts effect, unless value and probability are allowed to influence the processing of each other.

### Testing 10 resource-rational models in the ARRM framework

Next, we will show resource-rational models could provide a full explanation for the peanuts effect. Meanwhile, by comparing models with or without a specific resource-rational hypothesis, we can see whether the hypothesis holds for human decision behavior.

We constructed 10 resource-rational models in the ARRM framework, using three different base models with or without the RI or SP hypothesis, and fit the models to each participant’s 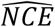 using maximum likelihood estimates (see Appendix and Supplemental Methods for details). The three base models were the Bounded Log-Odds model (BLO, Zhang et al., 2020), the Decision by Sampling model (DbS, Bhui & Gershman, 2018; Stewart et al., 2006), and the Rate-Distortion Model (RD, Bates et al., 2019; Gershman & Bhui, 2020), which have different resource-rational assumptions (see Methods). We used the Akaike Information Criterion (AIC; Akaike, 1974) as the metric of goodness-of-fit for model comparison. Unlike the Bayesian Information Criterion (BIC, Schwarz, 1978), AIC does not rely on the unrealistic assumption that the “true model” must be one of the models we tested (Gelman & Shalizi, 2013); AIC is also more robust than BIC for finite sample size (Vrieze, 2012). As shown in Figure 5A, the BLO+RI+SP model was the best-fitting model, achieving the lowest summed ΔAIC (the lower the better) in the JDA and JDB datasets. For participants collapsed across all the datasets, the group-level Bayesian model selection (Rigoux et al., 2014; Stephan et al., 2009) also shows that the BLO+RI+SP model outperformed the other models (higher protected exceedance probability in pairwise comparison).

**Figure 5.**
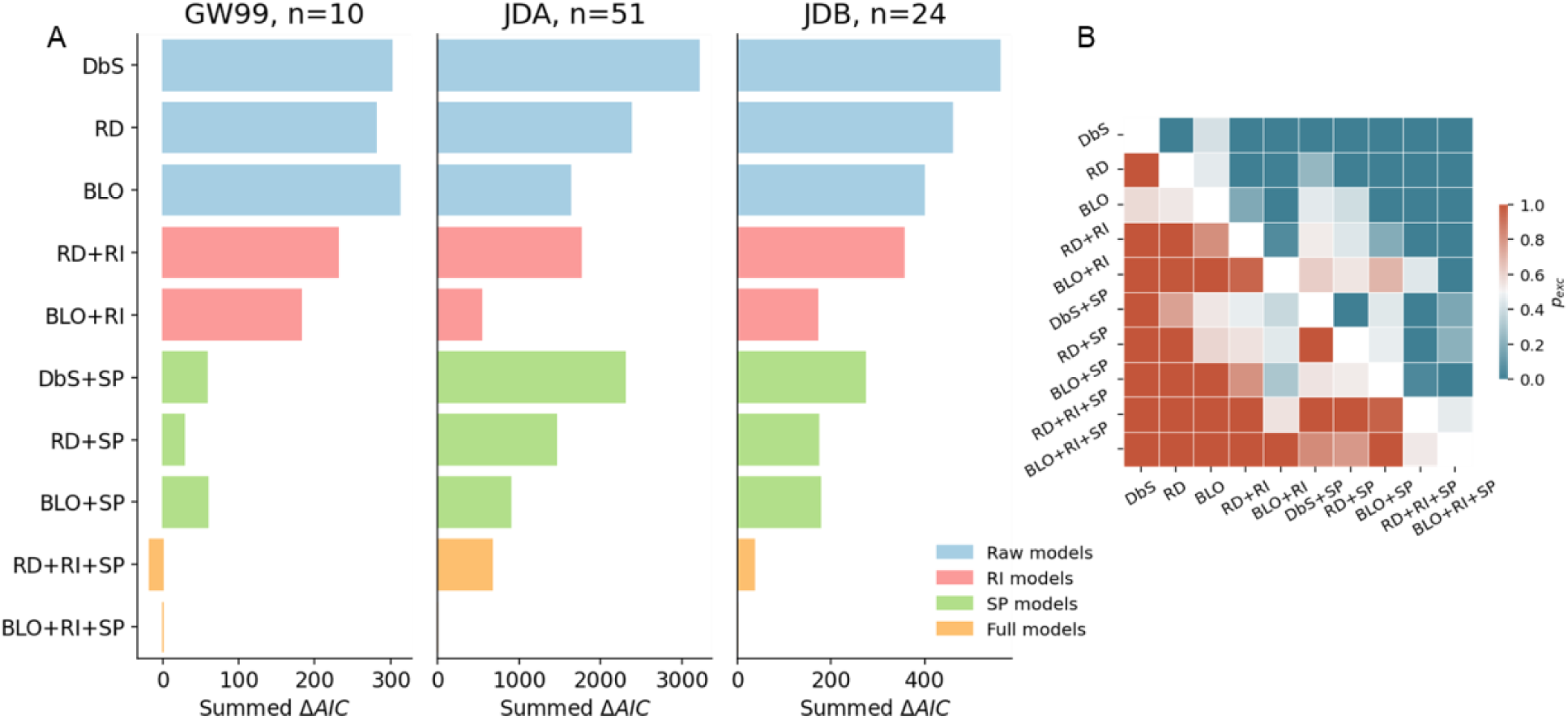
Model comparison results of the GW99 and ZRM20 (JDA and JDB) datasets. (**A**) Summed ΔAIC. The ΔAIC (with the BLO+RI+SP model as the reference) summed over participants is plotted as the metric of goodness-of-fit for each of the 10 models constructed in the ARRM framework. Smaller summed ΔAIC indicates better fit. To highlight the differences between models with and without the RI or SP hypothesis, models are color coded into four categories: base models (without RI or SP), RI models (with RI, without SP), SP models (with SP, without RI), and full models (with both RI and SP). Each panel is for one dataset or sub-dataset. In all datasets, the full models (orange) performed the best. (**B**) Results of group-level pairwise model comparison for participants collapsed across the GW99 and ZRM20 datasets. The protected exceedance probability (*P*_*exc*_, the probability that a specific model outperforms the other models), a group-level Bayesian measure of goodness-of-fit is calculated for comparison between each pair of the models. The darkest red and darkest blue respectively code that the row model outperforms the column model at a probability of 1 and 0.

By comparing models that only differ in one dimension of assumptions, we can see which assumption best agrees with human data. In most cases, models based on the coding scheme of BLO were better than those based on DbS or RD, though the AIC difference was moderate. Regardless of the choice of the base models (DbS, RD, or BLO), the models enhanced with the RI or SP hypothesis always fit better than those without them. The structural prior parameter *θ* (Eq. 4) in the winning model (BLO+RI+SP) was significantly greater than 0 (one-sample *t*-test, one-sided: *t*(84) = 2.506, *p* = 0.007), implying a prior belief consistent with the saying “the higher the risk, the greater the return”.

The BLO+RI+SP model can well capture both sub-effects of the peanuts effect as well as the inverted-*S*-shaped probability distortion (Figure 6). Intuitively, the sensitivity effect—the increase of sensitivity with gamble outcome—is explained by the RI hypothesis, which assumes that people would invest more cognitive resources to gambles of higher values. The elevation effect—the decrease of crossover point with gamble outcome—is explained by the SP hypothesis that assumes a negative correlation between probability and outcome value. According to the exchangeability test, the predictions of the BLO+RI+SP model show similar rejection of the separability assumption as the data do (Supplemental Figure S4).

**Figure 6.**
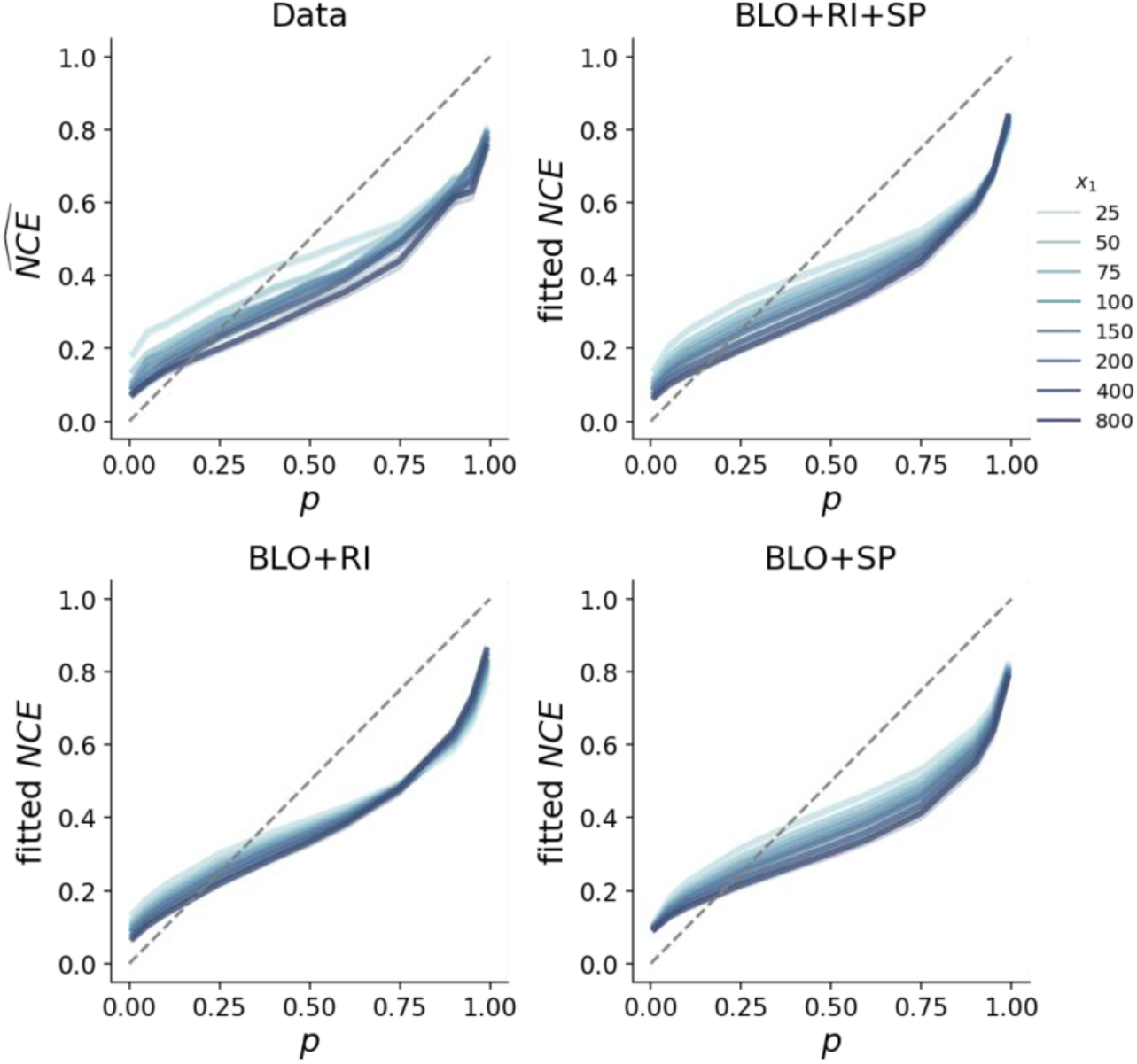
Data versus model predictions, pooled over the GW99 and ZRM20 datasets. Participants’ probability distortion curves (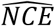, upper-left panel) versus model-predicted distortion curves (*NCE*, other three panels), separately visualized for gambles (*x*_1_, *p*; 0) with different *x*_1_. The prediction of the best-fit BLO+RI+SP model well captured both the sensitivity and elevation effect. Those of the BLO+RI and BLO+SP respectively captured the sensitivity and elevation effect, but not both, illustrating the necessity of both hypotheses. Conventions follow Figure 3.

To summarize, the BLO model enhanced by the RI and SP hypotheses provided a full explanation for the peanuts effect in decision under risk, with the hypotheses respectively accounting for the two sub-effects. Though the 10 models we considered above were all resource-rational models, a comprehensive set of non-resource-rational models were tested on the GW99 and ZRM20 (JDA and JDB) datasets by Zhang et al. (2020), according to which BLO outperformed all of a large set of decision models they tested, including the conventional CPT model and several heuristic-based models.. To further verify that decision theories based on other psychological mechanisms such as disappointment ( disappointment theory, Loomes & Sugden, 1986), regret (regret theory, Loomes & Sugden, 1982), or attention (salience theory, Bordalo et al., 2012) could not provide a better fit to the data, we constructed three additional models (see Appendix and Table 3 for details). Consistent with our reasoning earlier in the introduction of the peanuts effect, all three models fit much worse than our resource-rational models (Supplemental Figures S5 and S6).

**Table 3.**
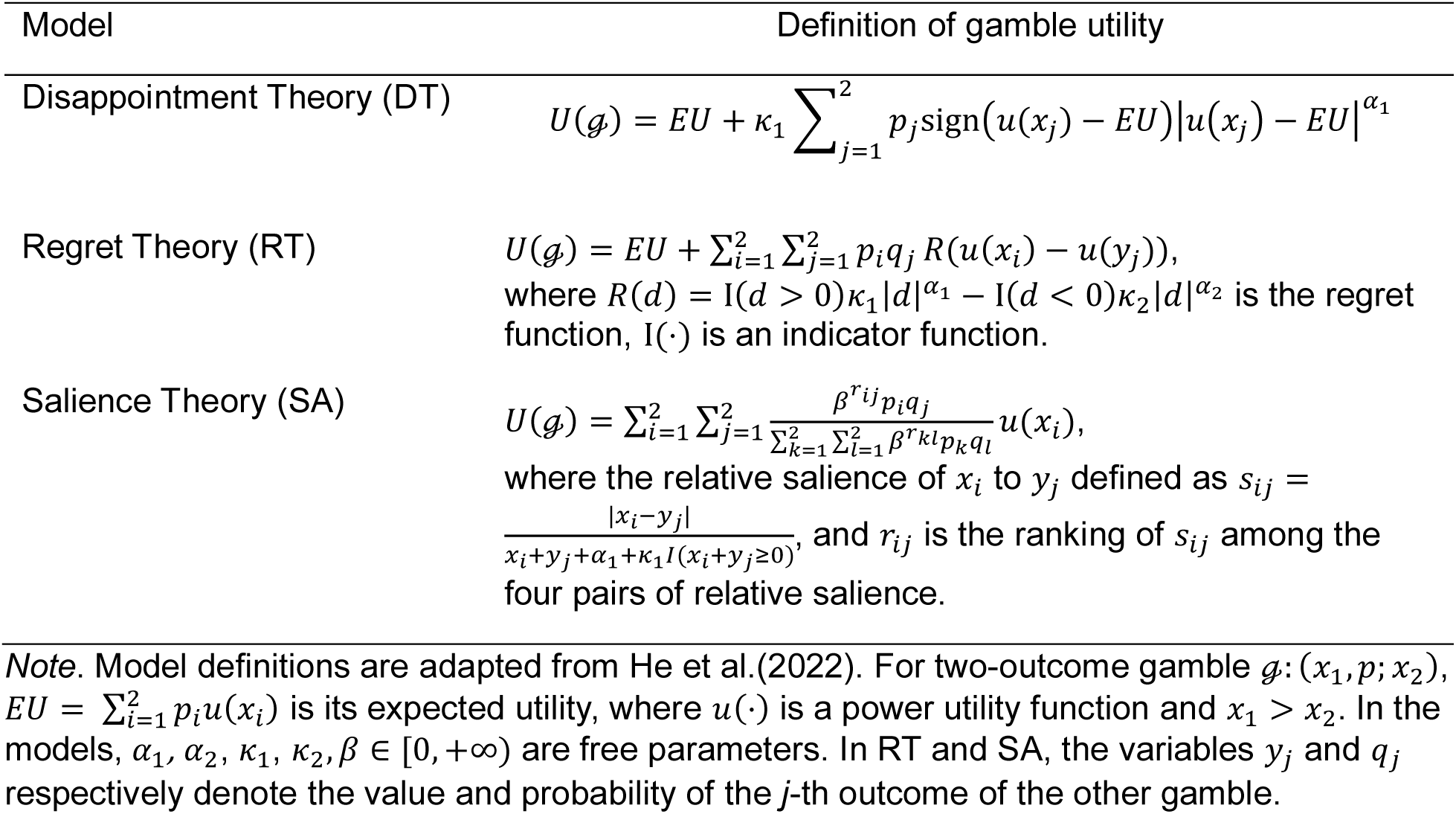
Models based on the disappointment theory, regret theory, and saliency theory.

## Study 2: replicating the peanuts effect and its ARRM modeling in a binary choice task

Both the GW99 and ZRM20 datasets we reanalyzed above were collected with the certainty equivalent task, where the options were presented in a tabular form, contrasting one gamble with a list of sure rewards in an ascending or descending order. This tabular presentation is efficient in eliciting certainty equivalents but is not the most typical scenario in decision making. To exclude the possibility that our findings only apply to the certainty equivalent task, we performed a new experiment of decision under risk using a binary choice task, where participants chose between a two-outcome gamble (*x*_1_, *p*; 0) and a sure reward on each trial. There were 8 levels of value *x*_1_ (25, 50, 75, 100, 150, 200, 400 or 800 Chinese Yuan) and 5 levels of winning probability *p* (0.05, 0.25, 0.5, 0.75 of 0.95), combining into 40 different gambles. Each gamble was repeated for 10 times, paired with different sure rewards that were selected by an adaptive procedure (see Methods in Appendix for details).

In this binary choice task (Figure 7AB, *n* = 21), we replicated the peanuts effect found in the certainty equivalent task of Study 1. According to the LMM analysis on participants’ 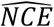, when the potential gain *D* of the gamble was greater, participants were more sensitive to the differences between different probabilities (*D* by probability interaction, *b* = 0.039, 95% CI [0.006, 0.071], *P* = 0.020), and had lower subjective estimates of the probabilities (main effect of *D*, *b* = −0.084, 95% CI [−0.128, −0.041], *P* < 0.001).

**Figure 7.**
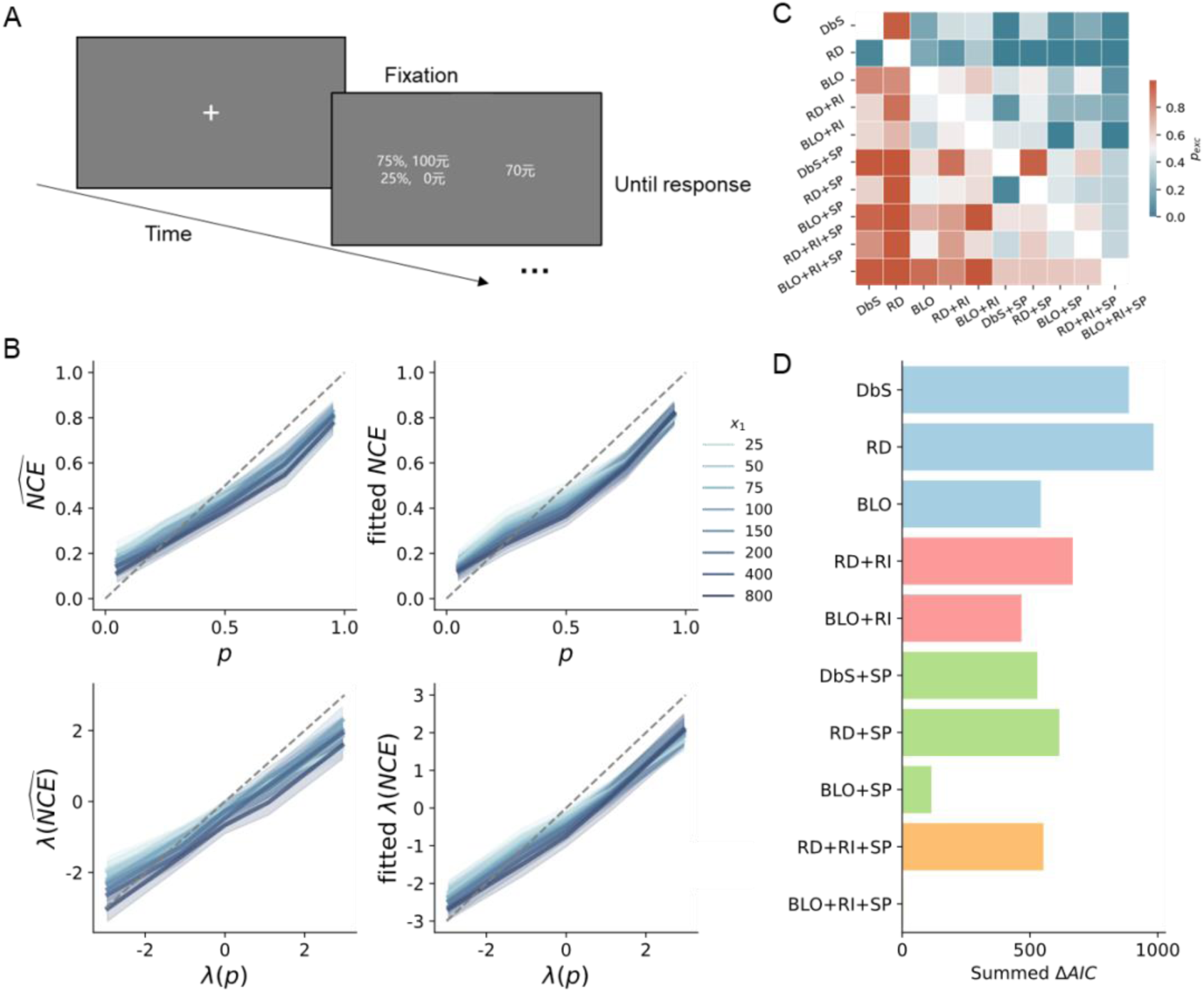
Task and results of the binary choice experiment (*n*=21). (**A**) The binary choice task. On each trial, participants chose whether to receive a gamble or a sure reward. The “元”in the display is the Chinese character for Yuan. (**B**) The measured peanuts effects (left) versus the predictions of the BLO+RI+SP model (right). To facilitate comparison with the results of the certainty equivalent task (Figure 3), participants’ choices were used to derive certainty equivalents, which were then transformed into normalized certainty equivalent (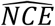, upper panels) and its log-odds (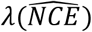, lower panels) for visualization. Conventions follow Figure 3. Similar to the results of the certainty equivalent task, gambles with a higher potential gain were associated with higher sensitivity and lower elevation in subjective probability (i.e., the two sub-effects of the peanuts effect). (**C**) Protected exceedance probability (*P*_*exc*_) for pairwise model comparisons. The darkest red and darkest blue respectively code that the row model outperforms the column model at a probability of 1 and 0. (**D**) Model comparison results. Conventions follow Figure 5. Again, the BLO+RI+SP model outperformed all other models in goodness-of-fit.

Similar to Study 1, the BLO+RI+SP model was the best model among the 10 resource-rational models in fitting participants’ binary choices (Figure 7CD). Again, the BLO base (coding scheme) outperformed the other two, and the models with the RI or SP hypothesis outperformed those without them, other things being the same. The structural prior parameter *θ* in the winning model (BLO+RI+SP) was again significantly greater than 0 (one-sample *t*-test, one-sided: *t*(19) = 3.597, *p* < 0.001). An additional LMM analysis on decision variability provides further support for the RI hypothesis (see Supplemental Figure S11 and its legend for details). Similar to Study 1, all the 10 resource-rational models outperformed the models based on the disappointment theory, regret theory, or salience theory (Supplemental Figures S7 and S8).

## Study 3: using ARRM to measure probability distortions under gain and loss

We have shown that the peanuts effect in decision under risk can be fully explained by resource-rational models developed in the ARRM framework, as the consequence of allocating more resources to higher-valued gambles (the RI hypothesis) and compensating for the covariation of value and probability in life experience (the SP hypothesis). But the datasets we analyzed in Studies 1 and 2 were limited to the gain domain (i.e., participants chose between risky and sure rewards). In Study 3, we extend our findings to the loss domain, reanalyzing the FD05 dataset from Fehr-Duda et al. (2010), where 153 participants were tested on both gain gambles and loss gambles. The dataset was also among the few decision studies that involved “real money” for participants instead of imaginary or small monetary payoffs.

As we introduced earlier, Fehr-Duda et al. (2010) were the first to propose that the peanuts effect reflects the effect of outcome value on probability distortion; they further measured the change of the probability distortion curve under different levels of values, separately for the gain and loss domains. In our reanalysis, we fit their dataset using the winning ARRM model, the BLO model enhanced with the RI and SP hypotheses. On one hand, we could see how well the model can explain probability distortion patterns in the loss domain. On the other hand, the model parameters would allow us to understand why probability distortion may differ between gain and loss, that is, whether the allocation of cognitive resources or compensation for prior beliefs may differ between the two domains.

Similar to Studies 1 and 2, we visualized probability distortions separately for gambles with different outcomes (Figure 8B) and ran an LMM analysis to verify the possible main and interaction effects of potential gain (*D*), probability, and outcome domain (gain or loss) on participants’ 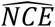. As to the elevation effect, participants’ subjective estimates of the probabilities decreased with the potential gain *D* (*b* = −0.141, 95% CI [−0.165, −0.117], *P* < 0.001), with an attenuated decreasing trend in the loss domain (*D* by IsLoss interaction, *b* = 0.121, 95% CI [0.090, 0.151], *P* < 0.001). Though participants’ sensitivity to probability did not change with *D* (*D* by probability interaction, *b* = −0.011, 95% CI [−0.028, 0.007], *P* = 0.239), a three-way interaction with the outcome domain (*D* by probability by IsLoss interaction, *b* = 0.048, 95% CI [0.023, 0.073], *P* < 0.001) indicates that the sensitivity effect might exist in the loss domain. The lack of sensitivity effect in the gain domain might reflect a lack of statistical power, because a significant sensitivity effect was identified in our following modeling analysis.

**Figure 8.**
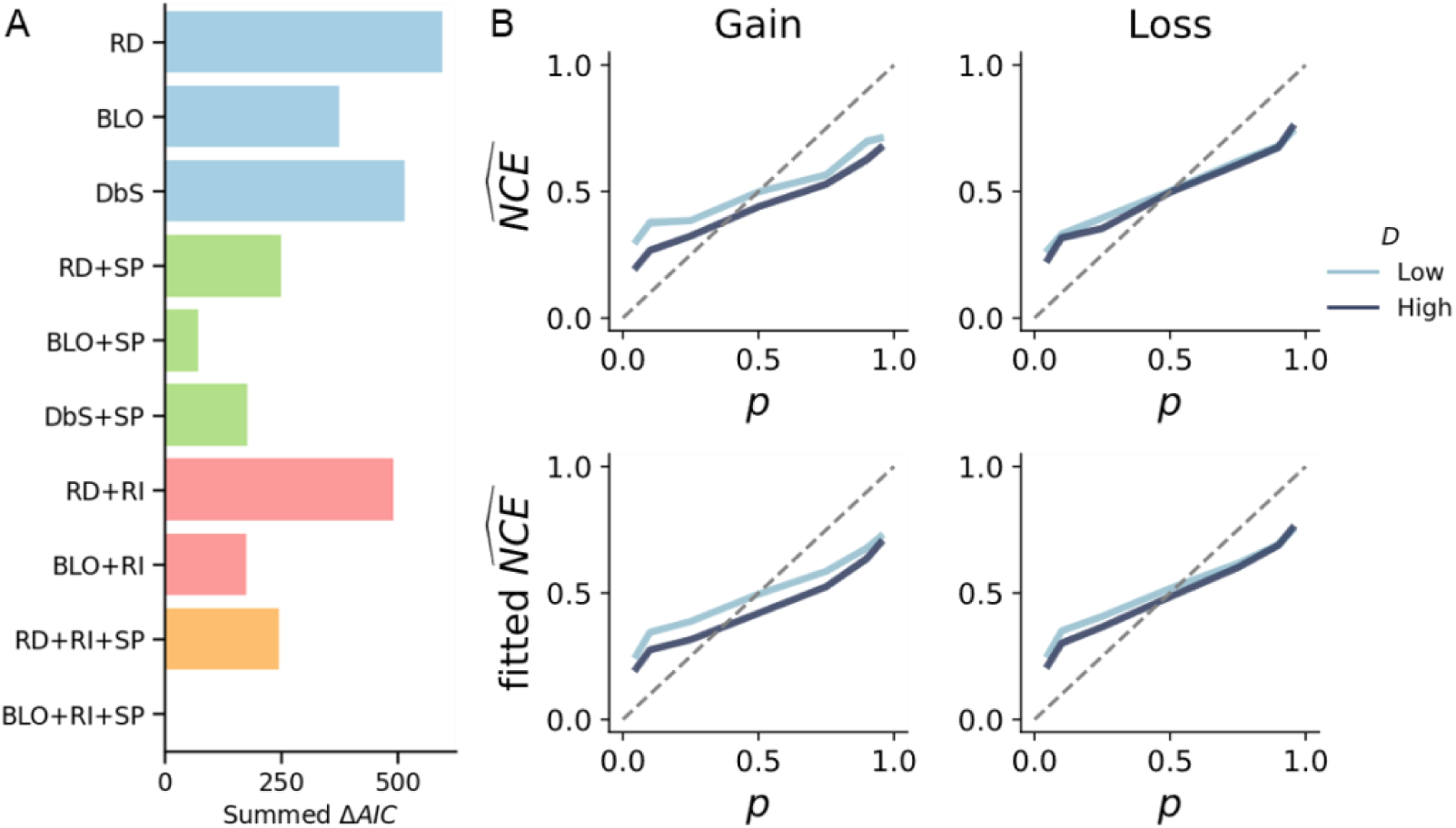
Modeling results for the FD05 Dataset (n=153). (**A**) Model comparison results for the FD05 dataset (153 participants, each of whom completed 56 trials). Different utility functions were used for gain and loss. Smaller ΔAIC indicates better fit. We did not (as for Studies 1 and 2) plot the results of group-level Bayesian model comparison, which shows negligible differences across the models. (**B**) Participants’ probability distortion curves (upper row) versus model-predicted distortion curves (lower row). The left and right columns are respectively for the gain and loss domains. Similar to Figure 6, participants’ normalized certainty equivalent 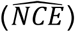 is plotted against objective probability (*p*). For visualization purposes, the gambles in the FD05 dataset were divided into two categories according to whether the potential gain *D* = |*x*_1_ − *x*_2_| of the gamble (*x*_1_, *p*; *x*_2_) was greater than 65 Yuan (darker curve) or not (lighter curve).

Again, according to the summed AIC, the best-fitting model to participants’ choices was the BLO+RI+SP model (Figure 8A), though group-level Bayesian model selection shows negligible differences between different models (the probability that the null hypothesis is true, BOR = 0.996). Resource-rational models still outperformed the disappointment, regret, or salience theory models (Supplemental Figures S9 and S10). To obtain reliable parameter estimates for further analysis, we also performed a hierarchical Bayesian fit for the BLO+RI+SP model. The fitted model can well capture participants’ behaviors, including the peanuts effect, in both the gain and loss domains (Figure 8B). Consist with our findings in Studies 1 and 2, participants tended to invest more cognitive resources to gambles involving larger stakes, with the group-level rational-inattention parameters (Eq. 2) in both the gain 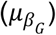 and loss 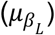 domains to be greater than 0 (95% HDI > 0; see Figure 9A, left panel). The group-level structural-prior parameters (Eq. 4) 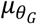 and 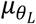 were also both greater than 0 (95% HDI > 0; see Figure 9B, left panel), as if participants held a prior belief that higher risk shall accompany higher gain or loss.

**Figure 9.**
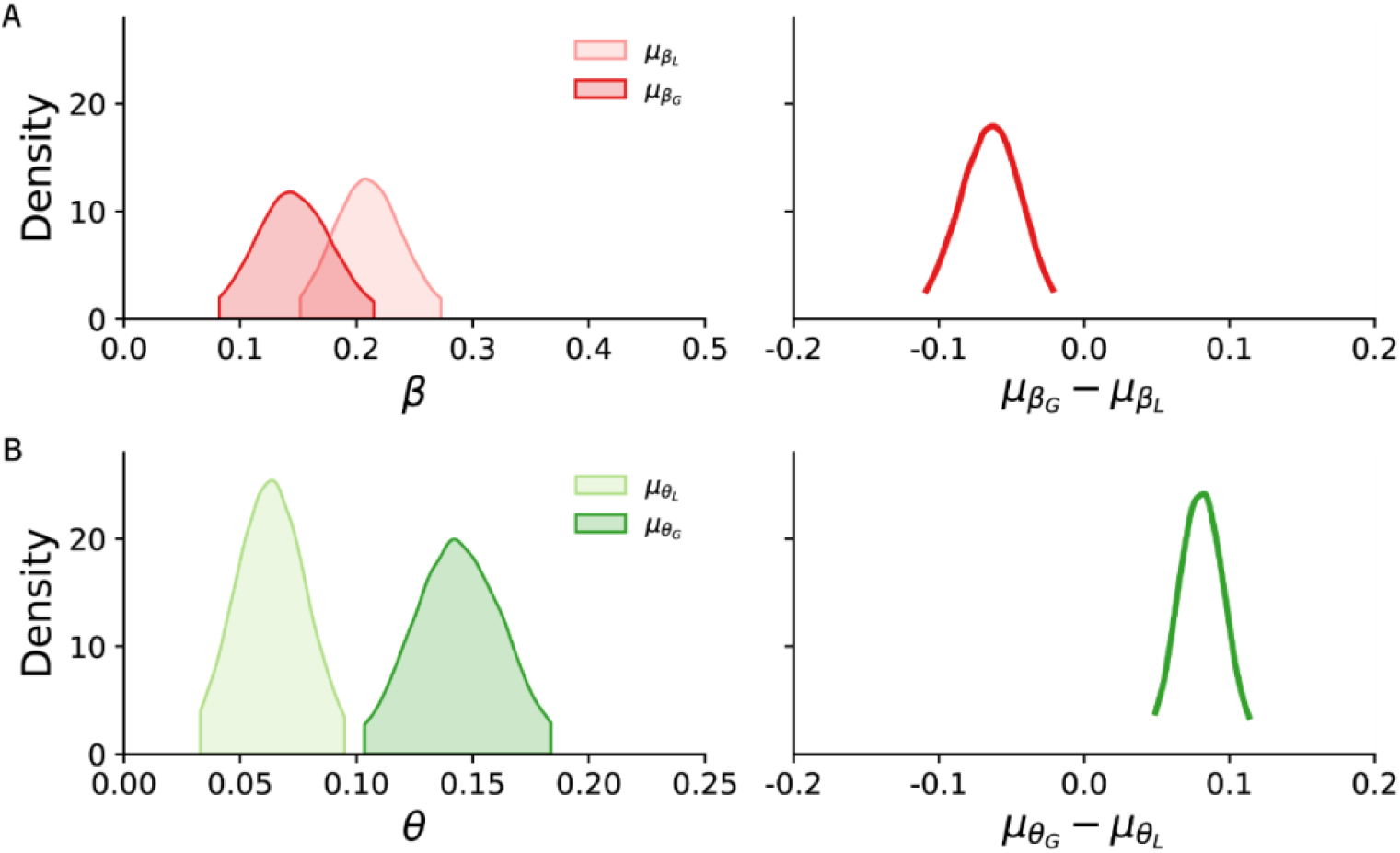
Gain versus loss: group-level rational inattention and structural prior parameters in FD05. The posterior distribution of each parameter (with 95% HDI) was estimated from hierarchical Bayesian modeling for the group of participants, separately for the gain and loss domains. (**A**) The posterior distribution of the rational inattention parameters 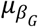, 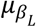 and their difference. Participants’ resource allocation was less sensitive to the change of outcome value for gain than for loss. (**B**) The posterior distribution of the structural prior parameters 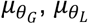 and their difference. Participants’ adjustment to the structural prior was more sensitive to the change of outcome value for gain than for loss.

The sizes of these effects were different for gain and loss gambles. Participants increased their resource investment less quickly for one unit of potential gain than for one unit of potential loss (the 95% HDI of 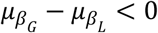, see Figure 9A, right panel). The extent to which participants compensated for a structural prior also differed between gain and loss, with a stronger negative relationship assumed for probability and gain than for probability and loss (the 95% HDI of 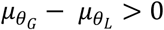, see Figure 9B, right panel).

## Discussion

Herbert Simon had a famous metaphor that the structure of the environment and the computational capabilities of the mind are the two blades of the scissors, together shaping human rational behavior (Simon, 1990). This idea of bounded rationality has motivated numerous studies on human heuristics and biases (Anderson, 1990; Lieder & Griffiths, 2020), but only recently been applied to modeling classic decision-making tasks such as decision under risk (Zhang et al., 2020) or intertemporal choice (Gershman & Bhui, 2020), tasks not traditionally perceived to be cognitively demanding.

Extending this approach, we developed ARRM as a computational framework to model rational agents who have limited cognitive resources to process the stimuli used for decision making but who use these resources appropriately to maximize task performance. In one new and three published datasets, we further demonstrated that ARRM can provide a full and normative explanation for human behavioral patterns in decision under risk, including a decision anomaly (the peanuts effect) that was difficult to be explained by previous decision theories. The seemingly inconsistency in human risk attitude turns out to reflect the allocation of more resources to more important probabilities and the compensation for the believed covariation of value and probability in the environment. The ARRM models can also serve as useful computational measures, providing insights into the differences in risk attitude between the gain and loss domains. The resource allocation is more sensitive to the change in loss than in gain, while the believed value-probability covariation is stronger in gain than in loss.

Below we discuss our work and its implications in the context of a broader range of decision theories and problems.

### Relationship of attention-dependent decision models to resource rationality

Some decision theories model human decision making as an attention-modulated valuation process, with the amount of attention allocated to a specific outcome determining its decision weight. For example, the transfer of attention exchange model assumes that the lower-valued outcomes in an option receive a larger proportion of attention than the higher-valued outcomes in the same option, when their probabilities are identical (Birnbaum, 2005). In contrast, the salience theory assumes that the same attributes of different options compete for attention in the decision process, during which higher-valued outcomes receive more attention due to their saliency and are thus endowed with higher decision weights (Bordalo et al., 2012; Mormann, 2016). Similarly, the parallel constraint satisfaction theory proposes that gambles with higher value and winning probability would lead to higher neural activation (Glöckner & Herbold, 2011). Further examples may include the decision field theory, which assumes decisions based on a dynamic sampling process, during which attention shifts among outcomes according to their associated probabilities (Busemeyer & Townsend, 1993; Johnson & Busemeyer, 2005).

Although attention is often viewed as a limited cognitive resource, these attention-dependent decision models are different from resource-rational models in two important ways. First, they often do not have normative motivations for why attention should be allocated in a particular way. As shown in the examples above, the attention allocation strategies assumed in different models often conflict with each other (Fiedler & Glöckner, 2012; Glöckner & Herbold, 2011; Johnson & Busemeyer, 2005). Second and more important, attention-dependent models often assume that attention directly influences decision weight, instead of influencing the precision of representation or information capacity as in resource-rational models. With more attention for larger gambles, attention-dependent models would not explain the peanuts effect but rather counterfactually predict that people would be more risk-seeking for larger gambles.

In our study, we found that the saliency theory model fit much worse than the resource-rational models to human decision behaviors. Other empirical findings are also more consistent with the assumption of our models that attention may influence the representational precisions of probability and value. On the individual level (Pachur et al., 2018), individuals who devote a higher proportion of attention to the probabilities of gambles have a steeper slope of probability distortion, implying more precise representation of probability, whereas those who allocate higher attention to the outcomes of gambles have higher sensitivity to the differences in outcomes. On the option level (Zilker & Pachur, 2021), people have a shallower slope of distortion for the probability in the risky option when they spend a larger proportion of time fixating the safe option.

The resource-rational and attention-dependent models do not necessarily conflict with each other but may reflect different levels of explanations. Recent analyses of eye-tracking and mouse-tracking data (He & Bhatia, 2023) suggest a resource-rational-like allocation of attention in decision under risk: Attention to the probability of a gamble is more likely when the previously attended gamble value is higher. If attention is assumed to influence the precision of representation instead of directly controlling decision weights as in the salience theory, attention-dependent models may serve as an algorithmic implementation of resource rationality. The relationship between attention and resource rationality may be elucidated in future experiments with eye-tracking or mouse-tracking recordings.

### Relationship of heuristic-based decision models to resource rationality

Heuristic-based models, which assume that human decision behaviors follow simple decision rules instead of maximizing expected utility, have been used to explain human decision biases that are unexplainable to CPT. Heuristic-based decisions often ignore part of the available information (Pachur et al., 2017). This can be viewed as a scenario where a resource-limited rational agent allocates resources exclusively to dimensions perceived as most crucial. For example, the similarity heuristic (Rubinstein, 1988) ignores the similar dimension of gamble options and focuses on their dissimilar dimension, which can be viewed as the extreme case of a rational cross-dimension resource allocation that invests more cognitive resources to the dimension with higher variance (C. J. Bates et al., 2019; Dewan, 2020). The more extreme maximax heuristic (Coombs et al., 1970) simply ignores the probability dimension and focuses on the option with the highest value, which yields suboptimal choices in general but might be rational when the contribution of probability to expected reward is small or the cognitive resource is extremely limited. Indeed, these heuristic-based decision algorithms are claimed to satisfy *ecological rationality* (Hertwig et al., 2022; Mata et al., 2012), according to which the heuristics are crystallized adaptive solutions of the brain to specific decision problems in the environment.

But unlike resource-rational solutions, these heuristic algorithms *per se* cannot vary with decision problems. In other words, each of them may approximate a resource-rational agent under specific resource constraints and task goals, but is not equivalent to the latter in general. Indeed, compared with heuristic-based decision models, resource-rational models can better explain human behaviors across different task environments (Callaway et al., 2022).

What were treated as heuristics in previous studies may enter the ARRM framework as prior beliefs. When people are found to judge the unknown probability of winning a larger reward to be smaller (Leuker et al., 2018; Pleskac & Hertwig, 2014) and respond faster in environments with negative correlations between outcome value and probability (Leuker et al., 2019), these behaviors were explained as a use of heuristics—higher risk often accompanies higher return. In our models, the same structural relationship is defined as a two-dimensional joint prior that contributes to a more precise estimation of probability. The prior, according to our taxonomy, belongs to a long-term prior that is likely learned through one’s life experience.

In our work, we did not directly compare the ARRM models with any specific heuristic-based models, but Zhang et al. (2020) have shown that the BLO model outperforms a representative set of heuristic-based models in fitting two of the datasets we reanalyzed here. Our best model (BLO+RI+SP), in turn, performed much better than the BLO model in fitting these datasets, and thus would outperform the set of heuristic-based models by transitivity.

### Biological plausibility of resource rationality

It is unclear whether or how an exact resource-rational agent may be implemented in the nervous system. As we argued earlier, approximate (or “bounded”) solutions to the resource allocation problems are more feasible than exact ones. For example, consistent with the hypothesis of rational inattention, larger incentives lead to higher arousal of the nervous system, which further modulates the encoding precision (Grujic et al., 2022; Mikhael et al., 2021).

Approximate resource-rational solutions may also be implemented by divisive normalization, a canonical neural computation originally found in the perceptual system (Carandini & Heeger, 2012) and later extended to higher-level neural circuits (e.g., valuation) as well (Louie et al., 2015). According to divisive normalization, the firing rate of a neuron to a specific stimulus is normalized by a population of neurons responding to similar stimuli, which allows neuron responses to adapt to the environmental statistics and achieve efficient coding (Simoncelli & Olshausen, 2001; Valerio & Navarro, 2003). In decision making, divisive normalization has been used to account for how the valuation of an option may change with the simultaneously (Louie et al., 2013) or recently (Khaw et al., 2017) presented options. Given that higher firing rate implies higher signal-to-noise ratio and thus higher representational precision, divisive normalization may be understood as allocation of cognitive resources (a total number of firing rates) among different options.

### Reconciling the different magnitude effects in different decision tasks

As we reviewed earlier, the magnitude effects in decision under temporal delay and decision under risk have seemingly opposite directions (Estle et al., 2006; Prelec & Loewenstein, 1991), if delay and risk are considered as parallel dimensions. However, we conjecture that these two types of magnitude effects arise from different cognitive algorithms. In our resource-rational model, the peanuts effect is treated as influences of value on probability distortion, partly arising from the prior belief “the higher the risk, the greater the return” (Leuker et al., 2018, 2019). Unlike us, Gershman and Bhui (2020) do not explain the magnitude effect as influences of value on temporal discounting. Instead, they argue that temporal discounting is an “as-if” phenomenon that originates from deteriorated precision of value under mental simulation of longer delay. Gershman and Bhui’s (2020) model for temporal discounting implies similar magnitude effects in different value domains, while our model for probability distortion implies domain-dependent peanut effects. Indeed, Jones and Oaksford (2011) found that the magnitude effect for cost is similar to their counterparts for gain or loss, but the peanuts effect for cost can be opposite to those of gain or loss.

### Other decision problems where ARRM may apply

When introducing the ARRM framework, we have summarized three different resource allocation problems and two different types of prior beliefs. These different modules, when assembled as needed, would allow us to provide normative explanations for a wider range of decision phenomena than previous resource-rational decision models did. Our modeling of the peanuts effect in decision under risk is just an example. Below we briefly discuss a few decision problems where ARRM may apply.

#### Decoy effects

One potential application concerns the decoy effects, that is, the probability of choosing one option against another may be dramatically changed by the presence of a third option (decoy) even when the decoy itself is rarely chosen (Louie et al., 2015). A memorizable example from Dan Ariely’s book “Predictably Irrational” (Ariely, 2010) is as follows. The reader of the journal *The Economist* was offered three subscription options: inexpensive e-copy, expensive hard-copy, or a bundle of hard-copy and e-copy of the same price as the hard-copy alone. Most people would choose the last option, though they would probably find the first option more attractive if the middle option did not exist. The decoy effects have diverse variants, depending on how the attributes of the decoy are related to those of the major options. Louie and colleagues (Louie et al., 2013, 2015) proposed that the decoy effects reflect context-dependent encoding of option values. From the perspective of resource rationality, the decoy effects may arise from the agent’s solutions to two resource allocation problems: how to allocate resources across options and across different attribute dimensions. These two problems can be interrelated; just as in our modeling of the peanuts effect, the value of one dimension (outcome) may influence the resource allocation on the other dimension (probability).

#### Decisions involving both delay and risk

The ARRM framework may also be applied to decision problems that involve both delay and risk, such as choosing among delayed gambles (e.g., 10% probability to receive $1M after 10 months). Including our models in the present article, the existing resource-rational models only apply to decision situations with delay or risk, but not both. When delay and risk are both involved, there turn out to be more sophisticated interactive patterns (Luckman et al., 2020; Vanderveldt et al., 2015, 2017), whose explanation is still elusive. The difficulty of modeling the effect, similar to the decoy effect discussed above, lies in the necessity to model the interrelated resource allocations across options and across attribute dimensions. ARRM can serve as a framework for developing such resource-rational models.

#### Limitations and Future Directions

Similar to existing resource-rational studies, we have only tested our models on tasks that are simple enough to be precisely predicted—choosing between a two-outcome gamble and a (or a series of) sure payoff(s). For gambles with more than two outcomes or tasks with multiple gambles, the resource-rational models are still to be worked out, due to the increased complexity of the resource allocation problems in more complex scenarios.

As a second limitation, we have only considered the influence of outcome value on the probability distortion process while keeping the utility function fixed, which follows the theoretical convention in the literature but is not necessarily the only possibility. Similar ARRM models may be constructed for the processing of value, with the value-probability interaction being the other way around: the decreased risk preference for larger outcome may be interpreted as increased sensitivity of value (i.e., greater exponent for the utility function) for larger probability. In which direction the interaction actually occurs will be a question for future studies.

A third limitation is the oversimplification in our modeling of noise. To make our resource-rational models computationally more tractable, we added a constant decision noise to each trial while neglecting the variation of representational noises across trials due to varied resource allocation. Moreover, we only considered the encoding noise in one dimension (probability) but neglected that of the other dimension (value). Though these are common practices in previous resource-rational models (Gershman & Bhui, 2020; Zhang et al., 2020), it would be valuable to develop more realistic models that include all sources of noises (Vieider, 2024) and test them using targeted experimental designs.

We assume people have limited cognitive resources and allocate their resources rationally for decision making. Though the behavior of such a resource-rational agent is derivable as the solution to an optimization problem, how to define the optimization problem is an empirical question. What are the cognitive resources, what are people’s prior beliefs for the environment, and how flexible the encoding and decoding schemes can be are all fundamental empirical questions that are still largely unknown. The module combinations tested in the present work are still limited. For example, for rate-distortion models, we chose the optimization problem to be minimizing squared error on the log-odds scale for analytical tractability, but did not know whether minimizing the squared error on the probability scale would better fit human data. The ARRM framework will enable us to develop new resource-rational models as well as to test them along with existing models to explore alternative theoretical possibilities. Further behavioral and neurophysiological measures to test these theoretical possibilities are also what the field calls for.

## Acknowledgments

HZ was supported by National Natural Science Foundation of China grants (32171095 and 32471152) and funding from Peking-Tsinghua Center for Life Sciences. Part of the analysis was performed on the High-performance Computing Platform of Peking University.

## Data and code availability statement

The data and codes that support the findings of this study are available in the Open Science Framework (OSF) repository upon publication of this article.

## Appendix

### Methods of Study 1

#### Experimental Design of the Datasets

Gonzalez & Wu’s dataset (GW99). The GW99 dataset (Gonzalez & Wu, 1999) includes the certainty equivalents (CEs) for 165 different virtual gambles from each of 10 US participants. In their experiment, each gamble was in the form of (*x*_1_, *p*; *x*_2_), that is, a probability of *p* to win $*x*_1_ and otherwise $*x*_2_, where *x*_1_ > *x*_2_ ≥ 0. There were 15 pairs of (*x*_1_, *x*_2_): (25, 0), (50, 0), (75, 0), (100, 0), (150,0), (200, 0), (400, 0), (800, 0), (50, 25), (75, 50), (100, 50), (150, 50), (150, 100), (200, 100) and (200, 150). These outcome pairs were combined with 11 probabilities (0.01, 0.05, 0.1, 0.25, 0.4, 0.5, 0.6, 0.75, 0.9, 0.95 and 0.99) to generate 15 × 11 = 165 gambles. For each gamble, the participant chose between the gamble and 12 different sure rewards (presented as two successive tables) in an adaptive two-stage procedure (see Gonzalez & Wu, 1999 for details), based on which the CE was estimated.

Zhang, Ren, & Maloney’s dataset (ZRM20: JDA and JDB). This decision dataset (Zhang et al., 2020) contains the CEs for 165 different virtual gambles from each of 75 Chinese participants from two sub-experiments (JDA: 51 participants; JDB: 24 participants). The JDA and JDB had almost identical procedures and designs as GW99, except that the outcomes were in Chinese Yuan instead of US dollars and participants were also tested on a separate task of relative frequency judgment (not analyzed here) in the experiment. The two sub-experiments differed from each other mainly in whether the trials of the decision and judgment tasks were interleaved (JDA) or in different blocks (JDB). Our purpose of presenting the two sub-experiments separately was to demonstrate the robustness of the results.

#### Statistical Analysis

For each participant and gamble, we calculated the normalized certainty equivalent 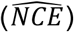 based on the measured CE, and then obtained its log-odds 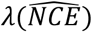. The 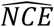 and 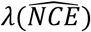 were used as nonparametric measures of the subjective probability. For each dataset, we performed the following linear mixed-effects model (LMM) analysis on 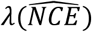 to test the potential influence of outcome values on probability distortion (as suggested by the peanuts effect):

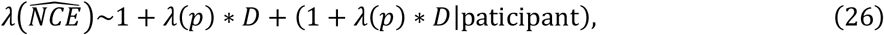

where *λ*(*p*) denotes the log-odds of the probability *p* in gamble (*x*_1_, *p*; *x*_2_), *D* = |x_1_ − x_2_| denotes the potential gain, 1 denotes the intercept term, *λ*(*p*) ∗ *D* denotes the main effects of *λ*(*p*) and *D* as well as their interaction, and (· |paticipant) denotes the random-effects terms. In the case the full model could not converge (which rarely occurred), we simplified the random effects structure following statistical guidelines (Barr et al., 2013; D. Bates et al., 2018; Brauer & Curtin, 2018). Data were normalized on the group level before entering LMMs. All the LMMs were implemented in python 3.8 with the statsmodels (v0.13.1) package and then replicated in R 4.0.3.

#### Model Fitting and Comparison

We fit the 10 models (described earlier) to the 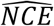 of each participant using maximum likelihood estimates. The Akaike Information Criterion (AIC; Akaike, 1974) was used to compare the goodness-of-fit among different models. We used the AIC of model BLO+RI+SP as a reference and calculated each model’s ΔAIC as its difference from the reference. Smaller ΔAIC indicates better fit. For each dataset or sub-dataset, we also performed the group-level Bayesian model selection (Rigoux et al., 2014; Stephan et al., 2009) and calculated the protected exceedance probability for each pair of models. We chose to use AIC instead of the Bayesian Information Criterion (BIC, Schwarz, 1978) as the major metric for model comparison, because BIC could measure how close a model comes to ground truth only when the “true model” is in the candidate set while AIC does not rely on such unrealistic assumption (Gelman & Shalizi, 2013). Besides, BIC can be less robust than AIC when the number of data points are relatively small (Vrieze, 2012). But for the reader’s reference, we still included the BIC results in the Supplement (Figures S6, S8, and S10).

#### The Exchangeability Test

The GW99 and ZRM20 datasets measured the CE for each gamble, this enabled us to conduct the model free exchangeability test. The measured CE curve for each gamble value *x*_1_ only has 11 probabilities, thus we first conduct a linear interpolation to approximate the full CE cure in probability range 0.01∼0.99. Given these CE curves, for any *x*_*a*_ > *x*_*b*_ > *x*_*c*_, we could find probabilities *p*_*b*_ and *p*_*c*_ so that the CE of gamble (*x*_*a*_, *p*_*b*_; 0) is *x*_*b*_ and the CE of gamble (*x*_*a*_, *p*_*c*_; 0) is *x*_*c*_. By applying Eq. 25, we have

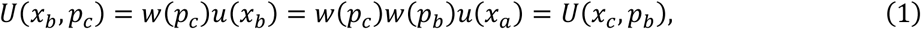

thus the gambles (*x*_*c*_, *p*_*b*_; 0) and (*x*_*b*_, *p*_*c*_; 0) should have the same CE.

For the GW99 and ZRM20 datasets, we draw 56 combinations from gamble values (*x*_1_, *x*_2_): (25, 0), (50, 0), (75, 0), (100, 0), (150,0), (200, 0), (400, 0), (800, 0). The value combinations [*x*_*a*_, *x*_*b*_, *x*_*c*_] were then passed into the interpolated CE curve of *x*_*a*_ to get the probability *p*_*b*_ and *p*_*c*_. Combinations with *p*_*b*_ and *p*_*c*_ outside the probability range 0.01∼0.99 (16 combinations) were excluded from analysis. The probabilities *p*_*b*_ and *p*_*c*_ were further passed into the CE curve of *x*_*c*_ and *x*_*b*_ respectively to get the measured 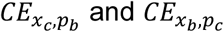. The measure CEs were scaled by *x*_*a*_ and then passed into paired t-test.

### Methods of Study 2

#### Participants

Twenty-one students (15 females, mean age = 20.6, SD = 2.2) in Peking University participated in our study. The experiment took approximately 1 hour and participants received 60 Chinese Yuan for their time. The study had been approved by the Committee for Protecting Human and Animal Subjects of School of Psychological and Cognitive Sciences at Peking University. All participants provided written informed consent and were paid for their time.

#### Design and Procedure

On each trial, a two-outcome gamble, denoted (*x*_1_, *p*; *x*_2_), and a sure reward were presented side by side on the screen. Participants need to choose between the two by pressing the “F” (left) or “J” (right) on the keyboard. The value of *x*_1_ had 8 levels: 25, 50, 75, 100, 150, 200, 400 and 800 (in Chinese Yuan). The value of *x*_2_ was always 0. The winning probability *p* had 5 levels: 0.05, 0.25, 0.5, 0.75 and 0.95. A combination of *x*_1_ with *p* led to 40 different gambles in total.

The sure reward paired with a specific gamble was selected according to the following two-round adaptive procedure. In the first round of 200 trials, the sure rewards for each gamble were set to 5 different levels, which were 0.1, 0.3, 0.5, 0.7 or 0.9 times of the *x*_1_ in the gamble. After the first round, for each gamble, the participant’s 5 choices were entered into a logistic regression against the sure-reward-to-*x*_1_ ratio to estimate a location parameter where the participant would be indifferent between the gamble and the sure reward (i.e., with a probability of 0.5 to choose sure reward). The location parameter was rounded to one decimal and added by −0.08, −0.04, 0, 0.4 or 0.08. The resulting ratio was then used to generate the 5 sure rewards for the gamble in the second round of 200 trials. In both rounds, trials for different gambles and sure rewards were interleaved in random order. For each participant, this two-round procedure was repeated twice, thus leading to 800 trials in total and 20 trials for each gamble.

#### Statistical Analysis

To make the results of this binary choice task comparable to those of the certainty equivalent task presented earlier, we estimated the certainty equivalent (CE) for each gamble using a logistic regression of choices against the sure rewards, based on all the 20 trials for the gamble. This estimated NCE 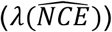 was used for visualization and further statistical analysis—an LMM similar to that used for the certainty equivalent task (Eq. 26).

#### Model Fitting and Comparison

Instead of fitting the estimated NCE, we fit the 10 models directly to participants’ binary choices (see the Decision Model section for details) using maximum likelihood estimates. As before, ΔAIC and protected exceedance probability were calculated for model comparison in goodness-of-fit.

The experiment design and statistical analyses were preregistered at OSF (https://osf.io/p76rg) for testing the peanuts effect. The modeling analysis (with the 10 models) were not preregistered^3^.

### Methods of Study 3

#### Fehr-Duda et al.’s dataset (FD05)

The experiment was performed in Beijing in 2005 on 153 Chinese participants, for each of whom the certainty equivalents for 56 different gambles (28 gain gambles and 28 loss gambles) were measured (Fehr-Duda et al., 2010). For each gamble, participants were asked to mark out their choices on a paper sheet where on the left side was the gamble and on the right side were 20 equally spaced sure payoffs. All gambles in FD05 had two outcomes, denoted (*x*_1_, *p*; *x*_2_) as before, where *x*_1_ > *x*_2_ ≥ 0 (gain gamble) or *x*_1_ < *x*_2_ ≤ 0 (loss gamble). The probability *p* had 7 levels: 0.05, 0.10, 0.25, 0.50, 0.75, 0.9, or 0.95. Half of the 28 gain gambles offered outcomes ranging from 4 to 55 Yuan (low-stake) and the other half offered 65 to 950 Yuan (high-stake). The 28 loss gambles had the same settings as the gain gambles except for reversing the sign of the outcomes. The highest gain or loss in the gambles amounted to several times of participants’ average monthly income at that time. Participants received monetary payoffs based on a random draw from their choices.

#### Statistical Analysis

Unlike the original paper (Fehr-Duda et al., 2010) that tested the effect of outcome value by dividing gambles into two categories according to the value of x_2_, we computed the potential gain (*D* = |x_1_ − x_2_|) for each gamble and used it as a continuous regressor. We used the following linear mixed-effects model (LMM) to examine the effects of outcome value and domain on the estimated NCE 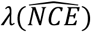, calculated as before):

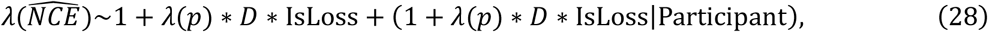

where (as before) *λ*(*p*) denotes the log-odds of the probability *p* in gamble (*x*_1_, *p*; *x*_2_), IsLoss denotes a dummy variable for outcome domain (0 for gain, 1 for loss), 1 denotes the intercept term, *λ*(*p*) ∗ *D* ∗ IsLoss denotes the main effects of *λ*(*p*), *D* and IsLoss as well as their interactions, and (· |paticipant) denotes the random-effects terms.

#### Hierarchical Bayesian Modeling

Because the gamble set for each participant and domain (28 gambles) was much smaller than that of Study 1 (165 gambles) or Study 2 (40 gambles), we fit the BLO+RI+SP model to participants’ NCEs using the hierarchical Bayesian method (Lee & Wagenmakers, 2013; Martin & Wiecki, 2018) instead of maximum likelihood estimates to obtain more reliable parameter estimates.

To extend our model to loss domain, we assumed that RI parameter *β* (i.e., the sensitivity of resource allocation to potential gain) and SP parameter *θ* (i.e., the sensitivity of compensating for the covariation between value and probability) could be different for gain and loss. We assumed each participant had two RI parameters (*β*_*G*_ and *β*_*L*_) and SP parameters (*θ*_*G*_ and *θ*_*L*_), with the subscripts *G* and *L* respectively denoting gain and loss. Following CPT, we assumed two different exponents *⍺*_*G*_ and *⍺*_*L*_ respectively for the power utility functions (Eq. 18) in gain and loss.

The model parameters were estimated using the Markov Chain Monte Carlo method (See Supplemental Text S2 and Table S4 for details), implemented by PyMC package on python 3.8. Four independent chains were run, each had 10000 samples after an initial burn-in of 5000 samples. The model had a good convergence with *R̂* ≤ 1.01 for all estimated parameters. The 95% highest density interval (HDI) was calculated for the group-level RI and SP parameters as well as their differences between the gain and loss domains.

### Alternative models tested outside resource rationality

Besides the 10 resource-rational models, we tested three models that represent the psychological mechanisms thought to be able to explain the peanuts effect: disappointment theory, regret theory, and salience theory. Each model enhances the computation of gamble utility with the influence of a specific psychological mechanism.

The disappointment theory (DT) assumes that participants’ choices are influenced by both the utility difference and the anticipated disappointment experienced when the realized outcome of a chosen gamble is worse than its expected outcome (Loomes & Sugden, 1986). The regret theory (RT) assumes that participants’ choices are affected by the potential regret they might feel if the outcome of the chosen option turns out to be worse than the outcome of the unchosen option (Loomes & Sugden, 1982). The salience theory proposes that the attention allocated to each outcome is proportional to its value, leading to an overweighting of the probability associated with the more salient (higher-value) outcome (Bordalo et al., 2012). We used the model forms summarized in He et al. (2022), with each model’s definition of gamble utility listed in Table 3. The definitions of their utility function and decision model are the same as those of the 10 resource-rational models.

## Footnotes

1 Markowitz (1952) constructed a parallel set of choices for the loss domain in a similar format as “X with certainty or take one chance in ten of owing Y?”. Contrary to the gain domain, most people seem to prefer the sure loss for small X and Y but the gamble for large X and Y. As we will see, the peanuts effect in both the gain and loss domains can be explained by the same model developed in the ARRM framework.

2 This equation holds when both outcomes are gains or losses, i.e., *x*_1_ > *x*_2_ ≥ 0 or *x*_1_ < *x*_2_ ≤ 0. When one is gain and the other is loss, they need two separate weighting functions.

3 When we preregistered the experiment, we had not known about the work by Fehr-Duda et al. (2010), who reasoned that the peanuts effect should be understood as the influence of outcome value on probability distortion. Instead, we had framed the peanuts effect as the influence of winning probability on the utility function, an alternative mathematical form to describe the peanuts effect (see Supplemental Text S1 and Figures S1 and S2). We later switched to the view of Fehr-Duda et al. (2010) to make a better connection to the decision literature.

