## Supplement for "Exploring the bounded rationality in human decision anomalies through an assemblable computational framework"

### Supplemental Text S1. An alternative mathematical form to describe the peanuts effect

When we preregistered the experiment, we had not known about the work by Fehr-Duda et al. (2010), who reasoned that the peanuts effect should be understood as the influence of outcome value on probability distortion. Instead, we had framed the peanuts effect as the influence of winning probability on the utility function, an alternative mathematical form to describe the peanuts effect.

In the cumulative prospect theory (CPT, as well as other decision models), the subjective utility of a gamble $\mathcal{g}: (x_1, p; x_2)$ is:

$$U(\mathcal{g}) = w(p)u(x_1) + \big(1 - w(p)\big)u(x_2), \tag{S1}$$

where $u(\cdot)$ and $w(\cdot)$ respectively denote the utility and probability weighting functions. Suppose the utility function $u(x) = x^{\alpha}$. In the special case that $x_2 = 0$, the certainty equivalent (CE) of the gamble follows:

$$CE^{\alpha} = w(p)x_1^{\alpha}. \tag{S2}$$

Taking logarithms and then scaling by $1/\alpha$ on both sides lead to:

$$\ln(CE) = \ln(x_1) + \frac{\ln(w(p))}{\alpha} \ , \tag{S3}$$

That is, regardless of the functional form of $w(\cdot)$, CPT predicts that the logarithm of $CE$ is a linear function of the logarithm of $x_1$, with slope equaling 1.

This form inspired us to use the following more general linear form (with slope parameter $k$ and intercept parameter $c$)

$$\ln(CE) = k\ln(x_1) + c \ , \tag{S4}$$

to measure the relationship between $\ln(CE)$ and $\ln(x_1)$ in the human data, separately for each specific probability $p$. If the estimated $k$ is not a constant but changes with $p$, the CPT prediction (Eq. S3) is violated, implying that the utility and probability weighting functions may not be independent of each other as assumed in CPT.

Indeed, by re-analyzing the measured CEs in published datasets (GW99 and ZRM20), we found a nice linear relationship between $\ln(CE)$ and $\ln(x_1)$ (for gambles with $x_2 = 0$), separately for each probability $p$ (Figure S1, left). However, the slope $k$ of the linear regression was less than 1, not constant but increased with $p$ (Figure S2, left).

We later performed a pre-registered experiment (Study 2 reported in the main text) to test whether there were similar effects in a binary choice task. We replicated the findings above as expected (Figures S1 and S2, right).

The linear regression based on Eq. S4 thus constitutes an alternative mathematical form to describe the peanuts effect. The resulting findings are consistent with those reported in the main text, indicating an interaction between the utility function and the probability weighting function. As we discussed in the main text, we still do not know whether it reflects the influence of probability on the utility function, or the influence of value on the probability weighting function.

**Figure S1**

*Relationship of CE to $x_1$ separately for different probabilities.*

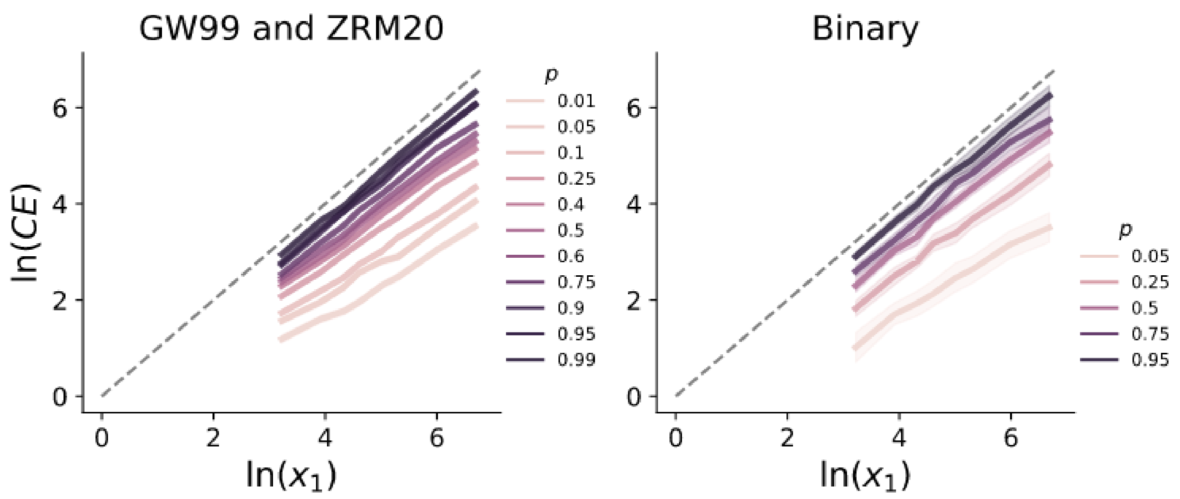


*Note*. Participants' CE for each gamble (with $x_2 = 0$) are plotted against the reward $x_1$ of the gamble on the logarithmic scale, separately for different winning probabilities (in different colors). Each curve is for one probability. Shadings denote SE across participants. Left: The GW99 and ZRM20 datasets of the certainty equivalent task (corresponding to Study 1 in the main text). Right: The binary choice task (corresponding to Study 2 in the main text).

**Figure S2**

*Regression slope $k$ of $\ln(x_1)$ as a function of the winning probability $p$.*

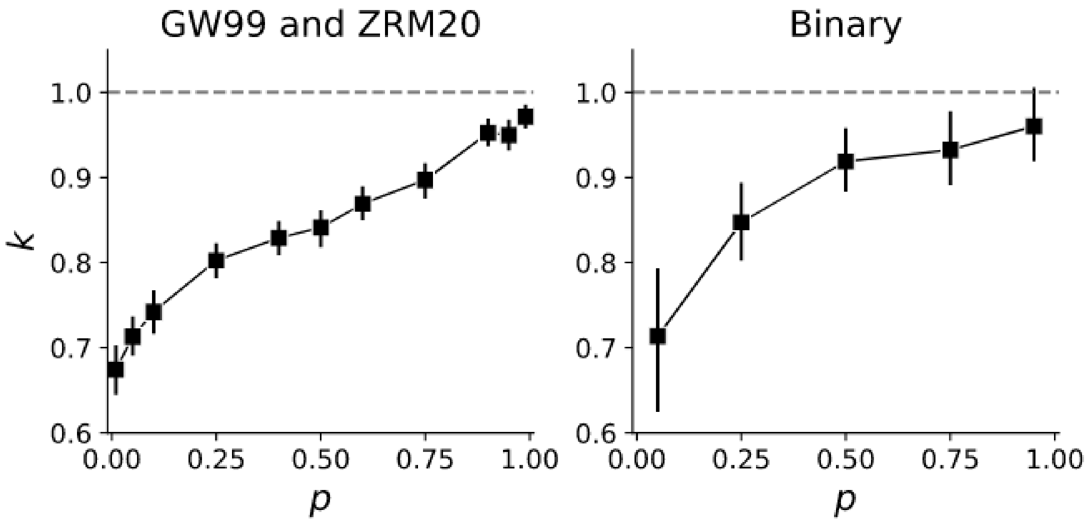


*Note*. Left: The GW99 and ZRM20 datasets of the certainty equivalent task (corresponding to Study 1 in the main text). Right: The binary choice task (corresponding to Study 2 in the main text). Error bars denote SE across participants.

### Supplemental Text S2. Hierarchical Bayesian model for FD05 dataset

We used a hierarchical Bayesian version of BLO+RI+SP model to fit the FD05 dataset (Fehr-Duda et al., 2010). We assumed that the rational inattention (RI) parameter $\beta$, structural prior (SP) parameter $\theta$ could be different in the gain and loss domains. Following the Prospect Theory, we assumed two different exponents $\alpha_G$ and $\alpha_L$ for the power utility functions in gain and loss. To boost the sampling process, we reparametrized the $\Delta^-$, $\Delta^+$ and $\Psi$ (Eq. 13) with $\mathrm{B}^-$, $\mathrm{B}$ and $\tau$:

$$\Delta^- = 3\mathrm{erf}(\mathrm{B}^-)\,, \tag{S5}$$

$$\Delta^+ = 3\mathrm{erf}(\mathrm{B}^- + \mathrm{B})\,, \tag{S6}$$

$$\tau = \frac{2\Psi}{\Delta^+ - \Delta^-} = \tau_0 + \beta_{G(or\ \mathrm{L})}\ln(|\mathrm{x}_1 - \mathrm{x}_2| + 1). \tag{S7}$$

where $\mathrm{erf}(.)$ is the Gaussian error function. Because the probabilities in the FD05 dataset were between 0.05 and 0.95, we bounded $\Delta^-$ and $\Delta^+$ between −3 and 3.

Parameters and prior distributions are listed in Table S1.

**Table S1**

*Parameters, priors and hyperpriors for the hierarchical Bayesian version of BLO+RI+SP model.*

| Parameters | Prior | Hyperpriors |
|---|---|---|
| $\kappa' = ln(\kappa)$ | $N(\mu_{\kappa'}, t_{\kappa'})$ | |
| $\mathrm{B}^-$ | $N(\mu_{\mathrm{B}^-}, t_{\mathrm{B}^-})$ | |
| $\mathrm{B}' = \ln(\mathrm{B})$ | $N(\mu_{\mathrm{B}'}, t_{\mathrm{B}'})$ | |
| $\lambda_0$ | $N(\mu_{\lambda_0}, t_{\lambda_0})$ | |
| $\tau_0{}' = ln(\tau_0)$ | $N(\mu_{\tau_0{}'}, t_{\tau_0{}'})$ | $\mu_{(\cdot)} \sim N(0, 1/9)$ |
| $\beta_L$ | $N(\mu_{\beta_L}, t_{\beta_L})$ | $t_{(\cdot)} \sim \Gamma(1, 1)$ |
| $\Delta\beta = \beta_G - \beta_L$ | $N(\mu_{\Delta\beta}, t_{\Delta\beta})$ | |
| $\theta_L$ | $N(\mu_{\theta_L}, t_{\theta_L})$ | |
| $\Delta\theta = \theta_G - \theta_L$ | $N(\mu_{\Delta\theta}, t_{\Delta\theta})$ | |
| $\alpha_G{}' = ln(\alpha_G)$ | $N(\mu_{\alpha_G{}'}, t_{\alpha_G{}'})$ | |
| $\alpha_L{}' = ln(\alpha_L)$ | $N(\mu_{\alpha_L{}'}, t_{\alpha_L{}'})$ | |

*Note*. $\mu_{(\cdot)}$ and $t_{(\cdot)}$ respectively denote the mean and precision of the Gaussian prior $N(\mu_{(\cdot)}, t_{(\cdot)})$ for each parameter.

### Supplemental Text S3. Simulation of rational resource allocation in decision under risk

According to the RI hypothesis implemented in our resource-rational models, the resource allocated to the probability $p$ in gamble $(x_1, p; x_2)$ increases linearly with the logarithm of its associated potential gain (Eq. 2). To verify that such resource allocation would indeed agree with that of a resource-rational agent, we performed a computational simulation in the experimental setting of Study 2, where on each trial participants chose between a two-outcome gamble $(x_1, p; 0)$ and a sure reward $c$.

For a resource-rational agent who has a limited cognitive resource to allocate among the different trials to represent probability and who aims to maximize the expected reward of the chosen option, the optimization problem can be reformulated as the following based on the Lagrangian duality principle:

$$J_p^{\text{optimal}}(x_1; \eta) = \underset{J_p}{\operatorname{argmin}}\, \mathbb{E}_{g_s}\left[Pr(C=0)c + \big(1 - Pr(C=0)\big)px_1 - \eta J_p(x_1)\right], \qquad (S8)$$

where $Pr(C = 0)$ is the probability of choosing sure reward (as specified in Eq. 34) and the terms $r(C = 0)c + \big(1 - Pr(C = 0)\big)px_1$ thus correspond to the expected reward of the agent's choice. The function $J_p(\cdot)$ of $x_1$ determines the resource allocated to the probability $p$ when the value of the gamble is $x_1$, whose logarithm roughly corresponds to the $R_g$ specified in Eq. 2. The parameter $\eta$ controls the balance between expected reward and cognitive cost (Mikhael et al., 2021; van den Berg & Ma, 2018). The $g_s$ is the distribution of gambles in the experiment.

In our simulation of the resource-rational agent, for each trial we could compute the decoded probability using the encoding and decoding scheme specified by Eq. 9, with the decoded log-odds $\widehat{\lambda_p} = \frac{J_p(x_1)\lambda(p) + J_0\lambda_s}{J_p(x_1) + J_0}$, where $J_0$ is a parameter of prior precision. Substituting the decoded probability into Eq. 19 and Eq. 24, we could obtain $Pr(C = 0)$. The decision temperature $b$, utility parameter $\alpha$, and prior mean $\lambda_s$ of the resource-rational agent were fixed at 1, 1, and 0, respectively. We searched for the optimal function $J_p(\cdot)$ for the optimization problem (Eq. S8) without constraining its functional form (i.e., allowing $J_p$ to be different for each different $x_1$).

The results of the simulation are shown in Figure S3. For any specific $\eta$, the optimal resource allocation implies that the resource allocated to the winning probability increases almost linearly with the logarithm of the potential gain, consistent with the assumption we used for resource-rational models (Eq. 2).

**Figure S3**

*Simulation results of the optimal resource allocation of a resource-rational agent*

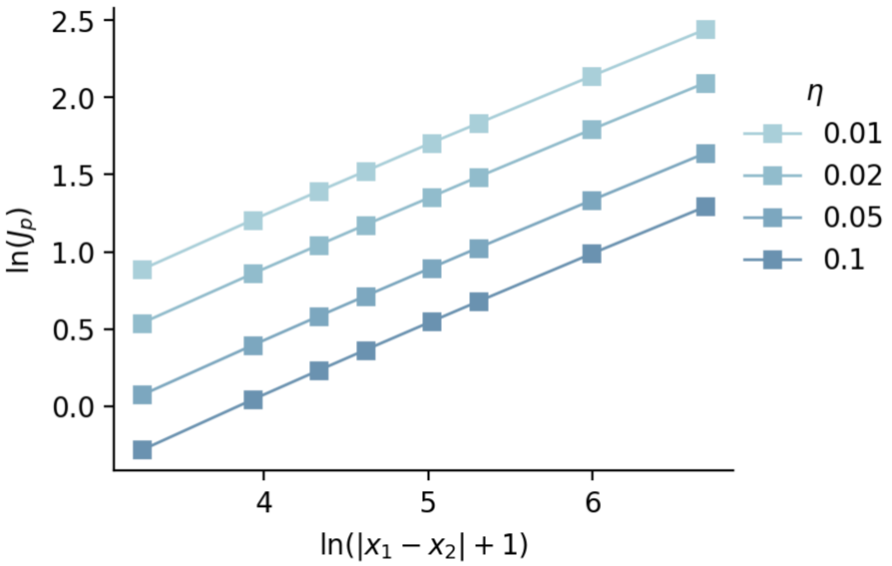


*Note*. Each line is for a specific level of sensitivity $\eta$ to cognitive cost. The decision temperature $b$, utility parameter $\alpha$, prior mean $\lambda_s$ of the simulated resource-rational agent were fixed at 1, 1, 0 respectively.

### Other Supplemental Figures

**Figure S4**

*The exchangeability test for the model predictions in the GW99 and ZRM20 datasets*

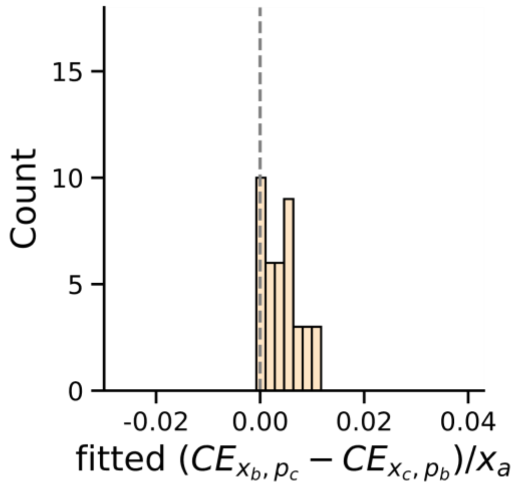


*Note*. Results of the exchangeability test: the histogram of the 40 equivalence pairs' CE differences. The exchangeability test was applied to the certainty equivalents predicted by the BLO+RI+SP model (mean prediction without noise). Similar to the results of the real data in Figure 4B, the mean fitted $(CE_{x_b,p_c} - CE_{x_b,p_c})/x_a$ of the 40 equivalence pairs were significantly greater than 0.

**Figure S5**

*AIC results of resource rational models versus three representative decision models for the GW99 and ZRM20 (JDA and JDB) datasets*

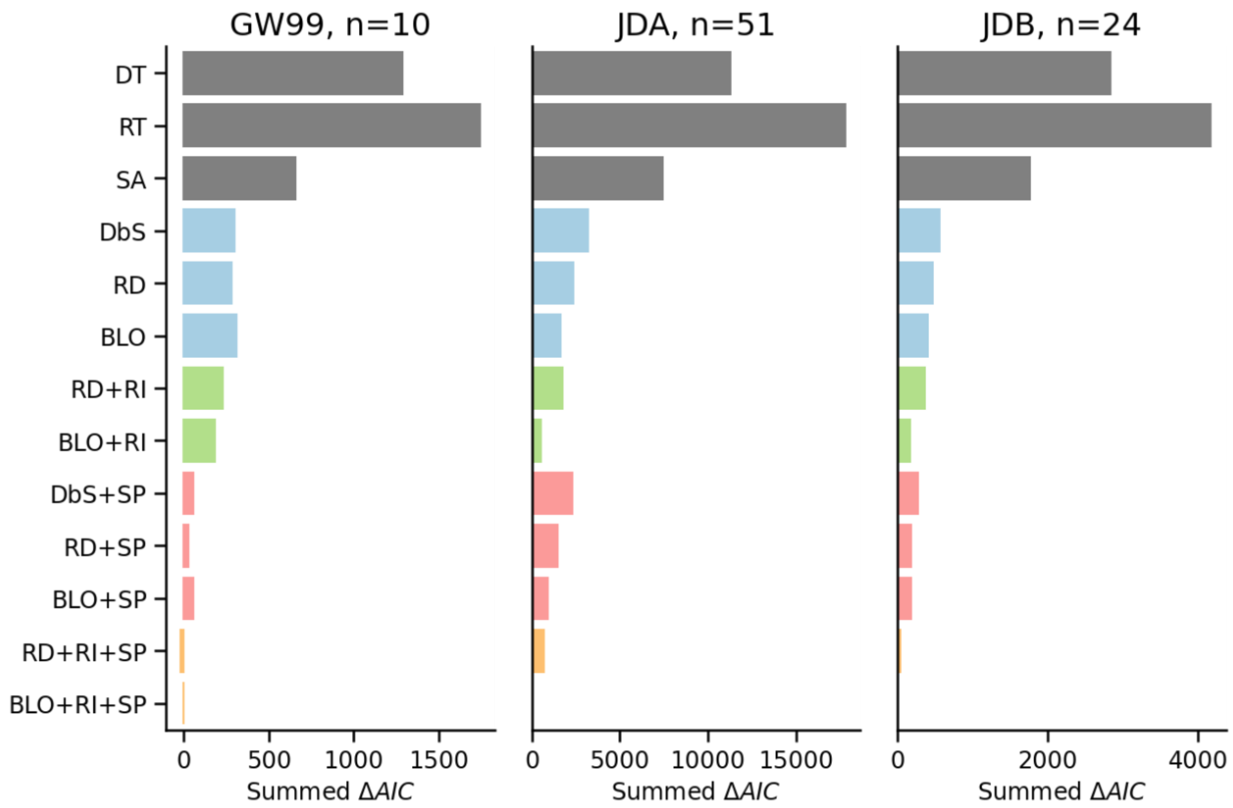


*Note*. The ΔAIC (with the BLO+RI+SP model as the reference) summed across participants is plotted for each of the 10 models constructed in the ARRM framework (colored bars) and three representative decision models outside resource rationality (gray bars). DT: Disappointment Theory. RT: Regret Theory. SA: Salience Theory. Smaller summed ΔAIC indicates better fit.

**Figure S6**

*BIC results of resource rational models versus three representative decision models for the GW99 and ZRM20 (JDA and JDB) datasets*

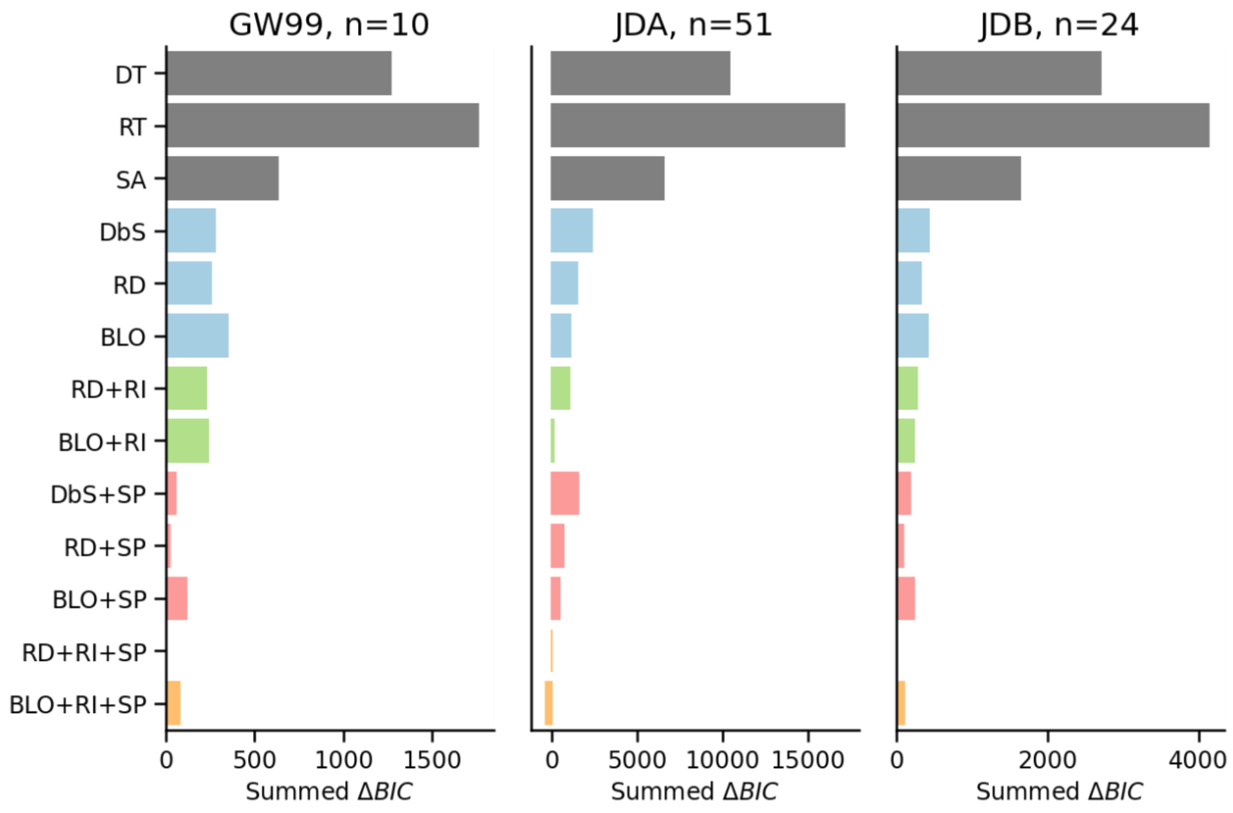


*Note*. The ΔBIC (with the RD+RI+SP model as the reference) summed across participants is plotted for each of the 10 models constructed in the ARRM framework (colored bars) and three representative decision models outside resource rationality (gray bars). DT: Disappointment Theory. RT: Regret Theory. SA: Salience Theory. The number of data points for each participant was 165. Smaller summed ΔBIC indicates better fit.

**Figure S7**

*AIC results of resource rational models versus three representative decision models for the binary choice experiment*

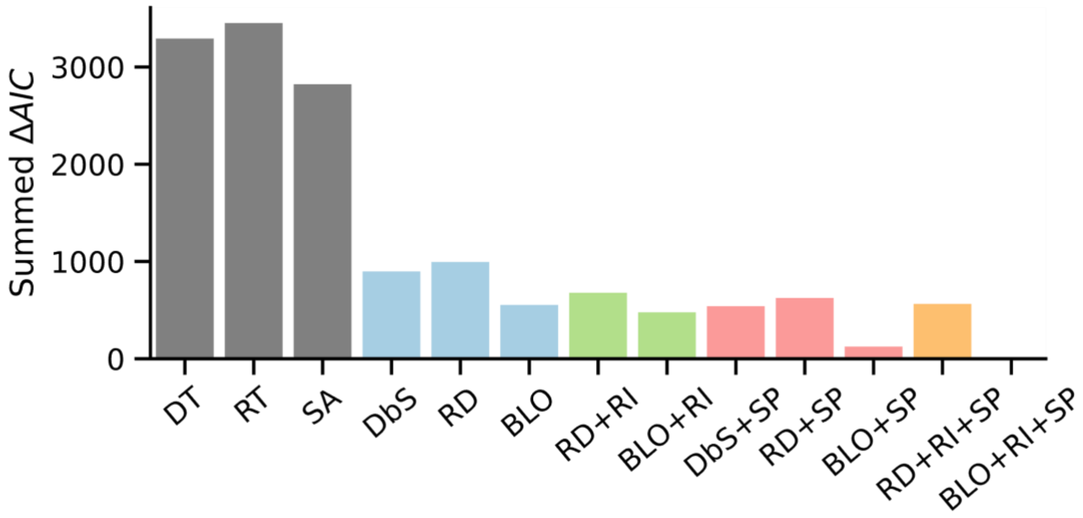


*Note*. The ΔAIC summed across participants is plotted for each of the 10 models constructed in the ARRM framework (colored bars) and three representative decision models outside resource rationality (gray bars). DT: Disappointment Theory. RT: Regret Theory. SA: Salience Theory. Smaller summed ΔAIC indicates better fit.

**Figure S8**

*BIC results of resource rational models versus three representative decision models for the binary choice experiment*

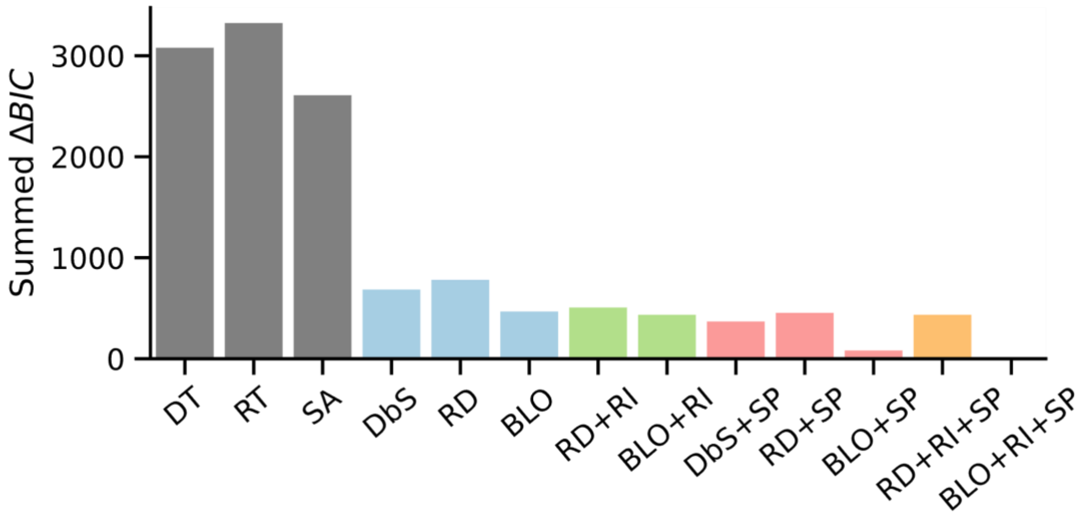


*Note*. The ΔBIC (with the BLO+RI+SP model as the reference) summed across participants is plotted for each of the 10 models constructed in the ARRM framework (colored bars) and three representative decision models outside resource rationality (gray bars). DT: Disappointment Theory. RT: Regret Theory. SA: Salience Theory. The number of data points for each participant was 800. Smaller summed ΔBIC indicates better fit.

**Figure S9**

*AIC results of resource rational models versus three representative decision models for the FD05 datasets*

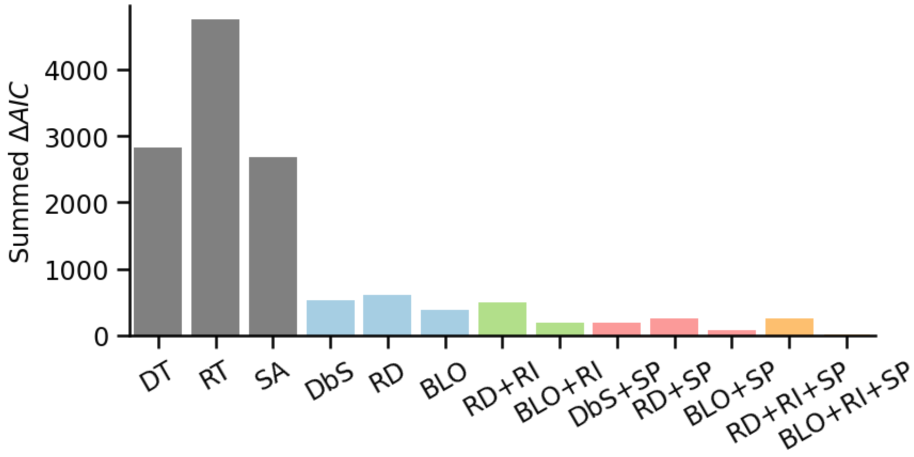


*Note*. The ΔAIC (with the BLO+RI+SP model as the reference) summed across participants is plotted for each of the 10 models constructed in the ARRM framework (colored bars) and three representative decision models outside resource rationality (gray bars). DT: Disappointment Theory. RT: Regret Theory. SA: Salience Theory. Smaller summed ΔAIC indicates better fit.

**Figure S10**

*BIC results of resource rational models versus three representative decision models for the FD05 datasets*

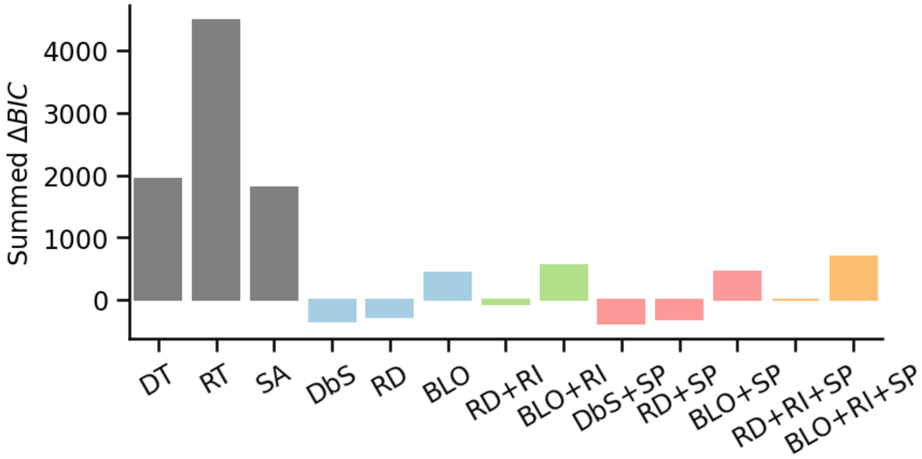


*Note*. The ΔBIC (with the RD+RI+SP model as the reference) summed across participants is plotted for each of the 10 models constructed in the ARRM framework (colored bars) and three representative decision models outside resource rationality (gray bars). DT: Disappointment Theory. RT: Regret Theory. SA: Salience Theory. Different utility functions are used for gain and loss. The number of data points for each participant was 56. Note that BIC may not be reliable when the number of data points is relatively small (Vrieze, 2012). Smaller summed ΔBIC indicates better fit.

**Figure S11**

*The change of decision noise with gamble value in the binary choice experiment*

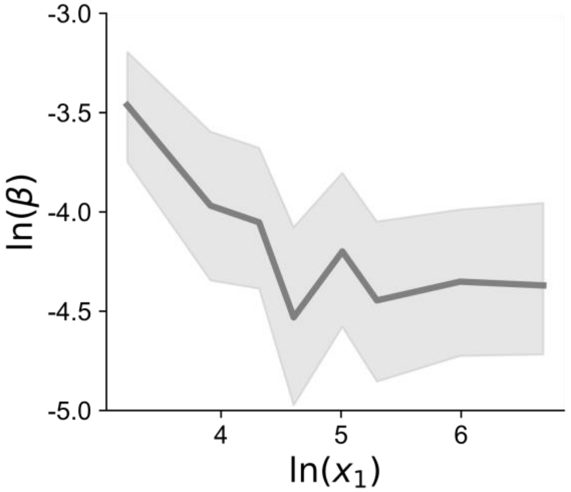


*Note.* The parameter $\beta$ plotted here indicates the temperature of the decision curve for a specific gamble, that is, how fast or slow the probability of choosing sure reward changes with the relative value of the sure reward. Higher temperature (shallower slope) implies higher decision noise. We performed a linear mixed-effects model on $\ln(\beta)$ with a random slope of $\ln(x_1)$ within participants and a random participant intercept to test the main effect of $\ln(x_1)$. The result indicates that across gambles the temperature $\beta$ decreases with increasing gamble value $x_1$ (*b* = −0.089, 95% CI [−0.171, −0.008], *P* = 0.031), which is consistent with our rational inattention assumption that more resources are used to represent the probability whose accompanying value is higher.
